# Dynamic optimization of extrachromosomal DNA copy number drives tumour evolution

**DOI:** 10.64898/2026.03.20.713026

**Authors:** Aditi Gnanasekar, Shu Zhang, Ellis J. Curtis, Jun Tang, Vishnu Shankar, Ivy Tsz-Lo Wong, Xiaowei Yan, Rui Li, Lu Yang, Melissa A. Yao, Andrea Ventura, Benjamin Werner, Howard Y. Chang, Paul S. Mischel

## Abstract

Extrachromosomal DNA (ecDNA) is common in human cancers and is associated with poor clinical outcomes, yet how ecDNA-driven genetic heterogeneity is translated into functional heterogeneity remains unclear. Using single-cell multiomics sequencing and multiplexed IF-FISH, we show that asymmetric inheritance of ecDNA generates copy number heterogeneity that propagates to gene expression programs, including oncogenic signaling and cellular stress responses. Transgenerational live-cell lineage tracking directly shows that ecDNA heterogeneity arises within only a few cell divisions and modulates daughter cell division timing in a copy number-dependent manner, a property not observed for evenly inherited chromosomal amplicons. We identify an optimal middle ecDNA copy number range that maximizes proliferative fitness at baseline, while drug selection pressure induced by low-dose CHK1 inhibition selects for cells with a new optimal range at low ecDNA copy numbers. These low ecDNA copy number cells pre-exist in the population and can be generated de novo, driving copy number shifts promoting drug resistance. In vivo experiments further demonstrate that shifts toward ecDNA copy numbers that are optimal under the tumour microenvironment enhance tumourigenicity. Together, these findings establish ecDNA copy number plasticity as a central driver of tumour evolution.

## INTRODUCTION

Extrachromosomal DNAs (ecDNAs) are circular DNA elements that are found in approximately 17% of human cancers^1^, more than half of all tumour types^2^, and are strongly associated with shortened patient survival^3^. EcDNAs are typically hundreds of kilobases to megabases in length^4–6^ and frequently encode oncogenes at high copy number^2,3,7–9^. Their circular architecture supports an open chromatin configuration that enhances chromatin accessibility and drives exceptional transcriptional output^4,10^. Furthermore, EcDNAs have been shown to cluster into transcriptionally active hubs that further amplify oncogenic signaling^11^. A defining hallmark of ecDNA is their asymmetric segregation during mitosis due to the lack of centromeres^12–14^, generating pronounced copy number heterogeneity within tumours that is associated with rapid adaptation and resistance to targeted therapies^2,14,15,16,17^.

Although ecDNAs are now recognized as active drivers of cancer evolution rather than passive byproducts of chromosomal instability, several key questions remain unresolved. It is still unclear what governs ecDNA abundance within individual cells, why cells do not continuously amplify ecDNA to maximize oncogenic potential, and why shifts in ecDNA copy number promote adaptation to different treatments or in different environments. Given the strong links between ecDNA copy number variability, cellular adaptation, and survival under drug treatment, clarifying how ecDNA dosage shapes cell fitness and fate is essential for improving our ability to predict and counteract ecDNA-driven cancer progression and drug resistance.

In this study, we investigated how ecDNA copy number status shapes single-cell behavior and phenotypic plasticity across ecDNA-positive cancers. We integrated single-cell multiomics analyses, live-cell imaging, computational modeling, and both in vitro and in vivo experimentation to define the functional consequences of increased oncogenic ecDNA copy number heterogeneity at single-cell resolution. Notably, we employed long-term live-cell imaging in an isogenic system to directly track the impacts of ecDNA abundance on cell survival and lineage dynamics in real time over multiple generations, representing the most direct assessment of how uneven inheritance of oncogenic amplifications influences cancer cell function and adaptation.

## RESULTS

### EcDNA copy number heterogeneity extends to heterogeneity in oncogenic pathways and cellular stress

Uneven inheritance of ecDNA to daughter cells during cell division generates immense amplicon copy number heterogeneity^14^. According to the central dogma, we hypothesized that such ecDNA copy number variation produces corresponding heterogeneity at the RNA and protein levels, driving variability in downstream pathway activity. We utilized two previously characterized^18^ near-isogenic cancer cell line pairs consisting a structurally similar amplicon encoded either on ecDNA or chromosomally as a homogeneously staining region (HSR), under an almost identical genetic background. Because HSRs undergo Mendelian inheritance, HSR(+) cells serve as an ideal control for dissecting the biological consequences of ecDNA’s uneven inheritance and the resulting copy number heterogeneity. DNA fluorescence in situ hybridization (FISH) performed in COLO 320DM and COLO 320HSR (carrying *MYC* amplicon), as well as GBM39-EC and GBM39-HSR (carrying *EGFRvIII* amplicon), confirmed that oncogene copy numbers in ecDNA(+) lines exhibit far wider distributions than oncogene copy numbers in HSR(+) cells (**Fig. 1a**). To provide an orthogonal sequencing-based method to profile cell-to-cell genetic heterogeneity, we applied a single-cell (sc) multiomics sequencing approach that jointly profiles scATAC-seq and scRNA-seq in ecDNA(+) and HSR(+) cancer cell lines (**Extended Data Fig. 1a**). Copy number was inferred from scATAC-seq using one-megabase windows, appropriate given the typical megabase-scale size of ecDNA^19^, and closely matched whole-genome sequencing estimates, indicating robust copy number readout with this analysis strategy (**Extended Data Fig. 1b**). Across multiple cancer cell lines, integrated scATAC-seq and scRNA-seq revealed that ecDNA(+) cells consistently showed strongly positive correlations between oncogene copy number and oncogenic transcript expression, and to a much greater extent than those of HSR(+) cells (**Fig. 1b, Extended Data Fig. 1c-d**). Housekeeping gene *ACTB* was not influenced by amplicon copy numbers, which confirmed that sequencing quality was not biased to copy number variance (**Extended Data Fig. 1c-e)**. Shannon indices, quantifying heterogeneity in expression of genes amplified on ecDNA or HSR and heterogeneity in hallmark pathway scores in COLO 320 and GBM39 isogenic cell line pairs, were generally increased in ecDNA(+) cells compared to HSR(+) cells (**Fig. 1c).** Delving more deeply into individual pathway scores, we found that cells with higher ecDNA copy numbers show elevated expression of downstream, cancer-driving pathways, including MYC-V2 targets in *MYC*-amplified ecDNA(+) cell lines and the PI3K/AKT-mTOR pathway in *EGFR* or *FGFR2*-amplified ecDNA(+) cell lines, compared to cells with lower ecDNA abundance (**Fig. 1d**). A greater level of oncogenic activity in the high copy number groups affirmed copy number’s impact on transcript expression extends to cellular and functional levels.

**Fig. 1.**
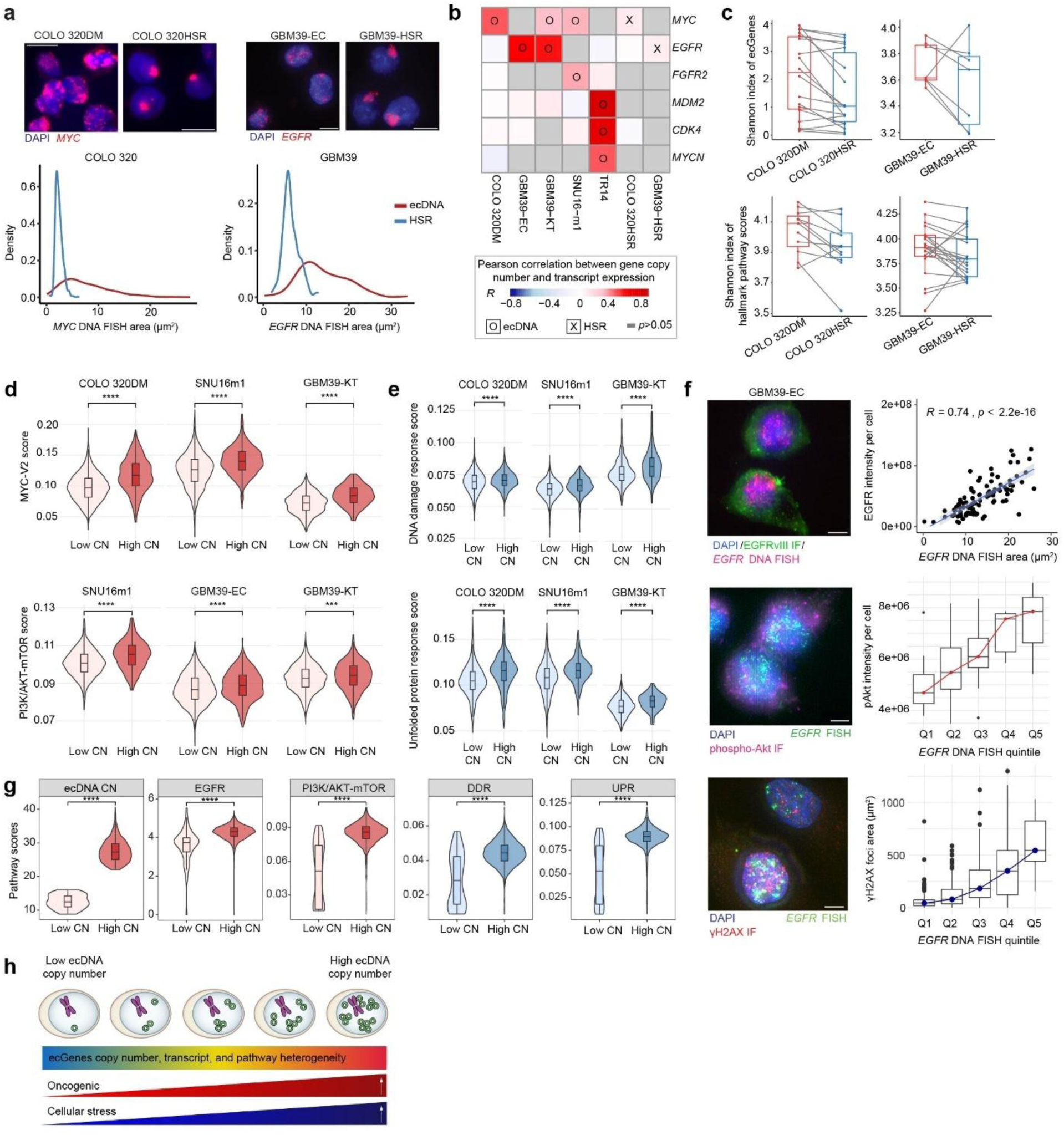
EcDNA-conferred copy number heterogeneity extends to transcript, protein, and pathway levels. **(a)** Representative images of DNA FISH for *MYC* and *EGFR* in COLO 320 and GBM39 isogenic cell line pairs, respectively (top). Scale bars, 10 µm. Quantification of ecDNA copy number measured by DNA FISH area for COLO 320 and GBM39 isogenic cell line pairs (bottom). **(b)** Heatmap depicting Pearson correlation coefficients between amplicon copy number and transcript expression from single-cell multiomics sequencing (scATAC-seq and scRNA-seq) in several cell lines. Strength of correlation (*R*) depicted only for statistically significant correlations (*p* < 0.05) **(c)** Shannon indices of the expression of genes amplified on ecDNA (top) and hallmark pathways (bottom) between ecDNA(+) and HSR(+) cells in COLO 320 (left) and GBM39 (right) isogenic cell line pairs. Wilcoxon rank-sum tests were performed to compare the mean gene and pathway expression between ecDNA(+) and HSR(+) cells. COLO 320DM vs. COLO 320HSR ecGene expression (p = 0.0016) and hallmark pathway scores (p = 0.024). GBM39-EC vs. GBM39-HSR ecGene expression (p = 0.22) and hallmark pathway scores (p=0.13). **(d)** Distributions of pathway expression scores for MYC-V2 targets (top) and PI3K/AKT-mTOR (bottom) expression between cells with high and low ecDNA copy numbers in cell lines with *MYC* ecDNA (top) and *EGFR* ecDNA (bottom). Statistical significance was assessed using a two-sided Wilcoxon rank-sum test. **(e)** Distributions of pathway expression scores for DNA damage response and unfolded protein response between cells with high and low ecDNA copy numbers in cell lines with *MYC* ecDNA (top) and *EGFR* ecDNA (bottom). Statistical significance was assessed using a two-sided Wilcoxon rank-sum test. (**f**) Representative images of combined FISH for *EGFR* ecDNA and IF for EGFR protein, phospho-Akt S473, and γH2AX (left). Scale bars, 5 µm. Pearson correlation scatterplot between *EGFR* copy number (measured by *EGFR* DNA FISH area) and EGFR protein expression (measured by EGFR intensity); n=97 cells (top right). Distribution of phospho-Akt Ser473 expression (measured by pAkt intensity); n=56 cells (middle right) and γH2AX expression (measured by foci area); n=818 cells (bottom right) across *EGFR* ecDNA copy number quintiles. **(g)** Distributions of EGFR protein abundance and pathway expression scores between cells with high and low *EGFR* ecDNA copy numbers in a human patient glioblastoma sample. Statistical significance was assessed using a two-sided Wilcoxon rank-sum test. **(h)** Schematic summarizing impact of ecDNA copy numbers on cancer cell phenotypes.

Previous studies have shown that high oncogene expression leads to high levels of replication stress, DNA damage response (DDR), and unfolded protein response (UPR)^20–24^. Consistently, we found that cells with higher ecDNA copy numbers also showed higher scores for cellular stress pathways, including the DDR and UPR, than cells with lower ecDNA copy numbers (**Fig. 1e**). Positive correlations between ecDNA copy number and downstream oncogenic activity, as well as cellular stress pathways, persisted across different cell-cycle stages, validated by sequencing and multiplex immunofluorescence (IF)-DNA FISH approaches (**Extended Data Fig. 1f-g**). In COLO 320DM cells with high ecDNA copy numbers, genes amplified on the ecDNA amplicon, including *MYC, PVT1, CDX2, CCDC26, PCAT1, and LRATD2*, were expectedly among the most highly expressed. In addition, MYC target genes, as well as several DDR and UPR genes, were significantly upregulated in the high copy number group. (**Extended Data Fig. 1h, Extended Data Table 1**). These results suggest that ecDNA-encoded genes may play a significant role in upregulating other cancer-driving genes not themselves encoded on the ecDNA amplicon.

To further corroborate the above gene signatures are reflective of functional protein correlations in ecDNA(+) cells, we performed multiplexed immunofluorescence (IF)-DNA FISH on fixed GBM39-EC cells from culture to measure EGFR protein abundance along with *EGFR* ecDNA copy number. EcDNA copy number was strongly positively correlated with protein abundance (**Fig. 1f**). Combined IF for phospho-Akt expression and DNA FISH for *EGFR* copy number revealed that ecDNA copy number is positively correlated with phosphorylation and activation of Akt, a pathway that is downstream of EGFR and is highly implicated in tumourigenesis^25^ (**Fig. 1f**), supporting copy number scales with oncogenic signaling activity. Combined IF for γH2AX (marker for DNA double strand breaks) and *EGFR* DNA FISH revealed that ecDNA copy number is also positively correlated with DNA damage (**Fig. 1f**), confirming ecDNA copy number scales with cellular stress as exemplified with DNA damage load. Beyond the neurosphere culture model, we quantified the relationship between ecDNA copy number and pathway activity in a human glioblastoma specimen with *EGFR* amplified on ecDNA profiled with SPACE-seq, a spatial multiomics approach integrating spatial ATAC-seq and RNA-seq^26^. Our results from analyzing the glioblastoma patient sample corroborated the trends observed across several ecDNA(+) cell lines, in which cells with high oncogene copy numbers showed greater expression in downstream oncogenic activity and cellular stress pathways (**Fig. 1g, Extended Data Fig. 1e**).

Collectively, these data demonstrate that ecDNA(+) cells exhibit elevated genomic heterogeneity, which contributes to more variable transcription and translation of ecDNA-encoded oncogenes and genes related to cellular stressors (**Fig. 1h**). These observations prompted us to investigate how cells balance the expression of oncogenic pathways and stress arising from elevated ecDNA copy numbers, and whether ecDNA copy number directly shapes, or even constrains, the fitness landscape of cancer cells.

### Live-cell imaging captures rapid emergence of ecDNA copy number heterogeneity

To track ecDNA dynamics and cell fitness, we engineered a near-isogenic pair of cell lines that allows real-time and persistent visualization of ecDNA and HSRs in living cells (**Fig. 2a**). Using a similar amplicon labelling strategy^14^, we used CRISPR-Cas9 to knock in a *tetO* array (96x) into the intergenic region between *MYC* and *PCAT1* in an isogenic colorectal cancer cell line pair: COLO 320DM and COLO 320HSR. The COLO 320HSR line exhibits Mendelian inheritance of the *MYC* oncogene and thus serves as a control for the uneven *MYC* inheritance observed in COLO 320DM^18^. We then visualized *MYC* ecDNA and HSRs with the expression of a Tet repressor fused to an mNeonGreen fluorescent protein (TetR-mNeonGreen) that bound to the *tetO* array insertion (**Extended Data Fig. 2a**). We named the ecDNA cell line TG19 ec and the HSR cell line TG19 HSR, by abbreviating *MYC-tetO* **T**etR-mNeon**G**reen to “TG” and “19” corresponding to the single clone number. DNA FISH on metaphase spreads confirmed robust *tetO* labeling efficiency in both TG19 ec and TG19 HSR (**Extended Data Fig. 2b**). The relationship between *tetO* area and *MYC* area is significantly linear and positively correlated in both clones, suggesting that TetR-mNeonGreen signal is a fair proxy for *MYC* copy number (**Extended Data Fig. 2c**). Overlap fraction between *tetO* and *MYC* foci was above 0.5 in 70% of metaphase spreads with DNA FISH (**Extended Data Fig. 2d**). There was no significant correlation between MYC copy number and the fraction of overlapping tetO and MYC foci in TG19 ec, indicating that *tetO* targeting and insertion efficiency is not preferentially biased toward higher copy number ecDNA. (**Extended Data Fig. 2e**). Quantification of the percent amplicon inherited to daughter cells in TG19 ec (n = 60) and TG19 HSR (n = 30) live single cell divisions confirmed that the per-cell amount of amplicon in a population of ecDNA(+) cells exhibits a wide normal distribution, while the HSR(+) cells show a narrow distribution, aligning with copy number distributions measured from fixed-cell staining in COLO 320DM and COLO 320HSR daughter cells (**Extended Data Fig. 2f**). Together, these results validate TG19 ec and TG19 HSR cell lines as reliable models for tracking *MYC* copy number and amplicon inheritance in real-time.

**Fig. 2.**
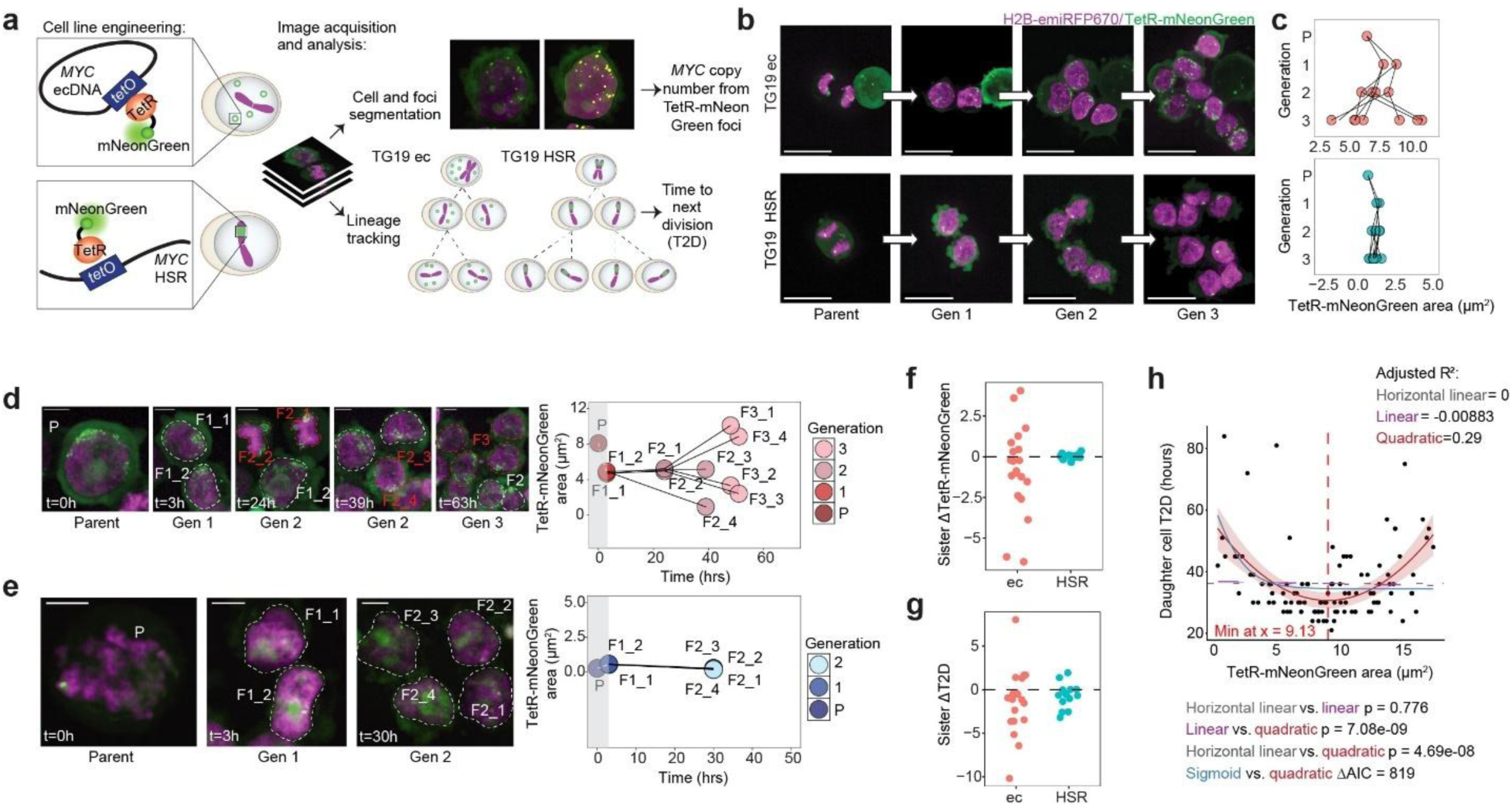
ecDNA copy number heterogeneity is generated rapidly and extends to division time heterogeneity. **(a)** Schematic of live-cell imaging cell line engineering and image acquisition and analysis pipeline. The engineered live-cell imaging cell lines originating from COLO 320DM and COLO 320HSR were named TG19 ec and TG19 HSR, respectively. **(b)** Representative images from live-cell imaging capturing a cancer cell as it divides and expands to a population of eight cells in TG19 ec (top) and TG19 HSR (bottom). Scale bars, 20 µm. **(c)** Quantification of TetR-mNeonGreen foci area in each cell in the TG19 ec and TG19 HSR lineages shown in (b). Each node represents a single cell, with connections relating parent cells to their daughter cells in the subsequent generation. TetR-mNeonGreen foci area was used as a proxy for *MYC* copy number. TetR-mNeonGreen foci area is likely underestimated in the parent generation due to condensed chromatin configuration at the time of image acquisition. **(d)** Representative images (left) and quantification (right) TetR-mNeonGreen foci area and time to division (T2D) from live-cell imaging and lineage tracking of TG19 ec. P = parent, F1_x = daughter cells (Generation 1), F2_x = granddaughter cells (Generation 2), F3 = great granddaughter cells (Generation 3). Red dashed outlines indicate the most recent division since the last timeframe. Scale bars, 5 µm. **(e)** Representative images (left) and quantification (right) of TetR-mNeonGreen foci area and T2D from live-cell imaging and lineage tracking of TG19 HSR. P = parent, F1_x = daughter cells (Generation 1), F2_x = granddaughter cells (Generation 2). Scale bars, 5 µm. **(f)** Quantification of TetR-mNeonGreen foci area difference between sister cells in TG19 ec (n=20 sister cell pairs) and TG19 HSR (n=18 sister cell pairs). Each dot represents the difference between two sister cells. **(g)** Quantification of T2D difference (hours) between sister cells in TG19 ec (n=24 sister cell pairs) and TG19 HSR (n=24 sister cell pairs). Each dot represents the difference between two sister cells. **(h)** Relationship between TetR-mNeonGreen foci area and T2D in TG19 ec cells (n=53 divisions, 106 cells). Horizontal linear, linear, sigmoidal, and quadratic models were applied to fit the data.

Live-cell imaging is a powerful method for tracking ecDNA in single cells as they divide over time, enabling simultaneous monitoring of ecDNA and cell dynamics. The isogenic live-cell system provides the most direct means for studying the accumulation of copy number heterogeneity over several generations and its putative impacts on cell fitness and fate (**Fig. 2a**). Imaging 60 to 80 hours enabled us to capture a single progenitor expanding to a population of 8 cells over three successive cell divisions (**Fig. 2b**). Hereafter, we refer to the single progenitor cell as parent, the two daughter cells as generation 1 (Gen 1), the four granddaughter cells as generation 2 (Gen 2), and the eight great granddaughter cells as generation 3 (Gen 3). The variance in proportion of *MYC* inherited in TG19 ec daughter cells steadily increased with each generation (TG19 ec lineage generation 3 variance = 8.676; coefficient of variation percent = 43%), while the proportion of *MYC* inherited in TG19 HSR daughter cells remained relatively constant from one generation to the next (TG19 HSR lineage generation 3 variance = 0.065; coefficient of variation percent = 24%) (**Fig. 2c**). This finding directly revealed that ecDNA-conferred copy number heterogeneity is generated rapidly, within a few cell divisions.

### Cell fitness is determined by inherited ecDNA content

In addition to quantifying *MYC* copy number, we also measured the time to the next division (T2D) as a proxy for fitness of each cell (**Fig. 2d-e, Extended Data Fig. 3a-b**). Most strikingly, TG19 ec sister cells arising from the same progenitor exhibited greater variance in both inherited *MYC* copy number and T2D compared to TG19 HSR sister cells (TG19 ec copy number variance = 7.20, T2D variance = 12.7 vs. TG19 HSR copy number variance = 0.0232, T2D variance = 2.02) (**Fig. 2f-g**). We only recorded T2D in cells from Generation 1 to 2 to avoid biases in division time arising from differences in cell cycle stage at the start of image acquisition or due to finite imaging duration. The relationship between ecDNA copy number and daughter cell T2D was best described by a quadratic model (**Fig. 2h**). This conclusion was supported by comparisons with nested linear models (horizontal vs. linear vs. quadratic) using ANOVA and non-nested nonlinear models (sigmoid vs. quadratic) using Akaike Information Criterion (AIC), where lower AIC values indicate a better model. The sigmoidal model, commonly used to capture dose-response relationships and here intended to model a plateau in T2D beyond a copy number threshold, provided a substantially poorer fit than the quadratic model (AIC_sigmoid = 813.7 vs. AIC_quadratic = −5.1). These analyses indicate that daughter cell fitness is maximized at a middle ecDNA copy number range (minimum near 9.13 µm^2^ of TetR-mNeonGreen area in this dataset) rather than increasing or saturating monotonically with ecDNA abundance. We therefore defined the ecDNA copy number range associated with the shortest T2D as the optimal copy number range for maximal proliferative fitness. Integration with single-cell multiomics analyses suggests that this optimal range reflects a balance between promoting oncogenic activity while minimizing cellular stress (**Fig. 1d-h**). Importantly, our live-cell system demonstrated that ecDNA abundance has an immediate effect on cancer cell fitness, measured by T2D.

There were a few outliers with unexpectedly long T2D despite having optimal copy numbers and outliers with unexpectedly short T2D despite having suboptimal copy numbers. Via retrospective lineage tracking, we were able to identify the parent cell and measure its copy number from TetR-mNeonGreen foci area. Interestingly, parents of cells with slow T2D had more extreme copy numbers, farther from the value of 9.13 µm^2^, while parents of cells with fast T2D had more optimal copy number, closer to the value of 9.13 µm^2^ (**Extended Data Fig. 4a**). These results suggest that the parent cell ecDNA copy number may also play a role in predicting daughter cell fitness, indicating a transgenerational effect across somatic cell divisions. Combined IF for Aurora B kinase (marker for paired daughter cells) and γH2AX with DNA FISH for *MYC* revealed greater variance in inherited damaged DNA in TG19 ec compared to TG19 HSR daughter cells (**Extended Data Fig. 4b**). In addition to taking longer time to divide, cells with high copy numbers generated ecDNA(+) micronuclei, which are extranuclear structures characterized by chromosomal instability and damaged DNA^27–30^. Based on the link between ecDNA copy number and DNA damage^18^, along with the relationship between ecDNA copy number and micronuclei^31^, we evaluated the association between ecDNA(+) micronuclei and cellular fate. Cells with large (> 2 µm diameter) ecDNA(+) micronuclei suffered worse fate (underwent cell death or remained undivided) with greater probability than their sister cells which had comparable ecDNA copy numbers but no or smaller micronuclei^31^ (**Extended Data Fig. 4c**). These results suggest that it is not merely the amount of ecDNA, but the integrity of the inherited ecDNA that impacts cell fitness. Since ecDNA copy numbers positively correlate with increased DNA damage (**Fig. 1d-g**), cells with high ecDNA copy numbers that manage to divide may pass on differential amounts of clustered and damaged ecDNA^31^, predisposing one daughter cell to a worse fate.

To verify the existence of optimal copy number across ecDNA(+) cancer cell lines, we combined IF for Aurora B kinase and Cyclin A with DNA FISH for *MYC* to evaluate differences between copy number of reproducing cells versus the copy number of all G1 cells (Cyclin A-negative) in the population (**Extended Data Fig. 5a**). We calculated the parent cell copy number by averaging the copy numbers in daughter cells and then compared parent cell copy number to the copy numbers of all G1 cells in the population. We specifically compared parent cells to G1 interphase cells to avoid bias towards higher ecDNA copy numbers resulting from DNA being replicated and genomic content being doubled in S and G2/M phase cells (that are positive for Cyclin A). In both COLO 320DM and PC3 DM, there is a higher peak at the second copy number quintile for parent cell copy number versus that of other G1 interphase cells, indicating that there exists an optimal copy number for proliferation that falls in a middle, narrower copy number range than the copy number range of the full population (**Extended Data Fig. 5b**). Together, these findings support the notion that there exists an optimal ecDNA copy number range associated with the best fitness.

### Dynamic optimal ecDNA copy numbers drive cell adaptation

To understand how the relationship between ecDNA copy number and T2D can inform cancer cell evolution, we first simulated population growth from a range of initial copy numbers using three models: (1) a horizontal model, in which T2D is independent of copy number (**Extended Data Fig. 6a**); (2) a sigmoid-plateau model, in which T2D decreases and then plateaus as copy number rises (**Extended Data Fig. 6b**); and (3) a quadratic model derived from live-cell imaging, in which T2D is minimized at a middle copy number range (**Extended Data Fig. 6c**). Under the quadratic model, the mean ecDNA copy number of the population rapidly converged to an optimal range as the population expanded (**Extended Data Fig. 6f**), whereas no convergence was observed under either the horizontal or sigmoid-plateau models (**Extended Data Fig. 6d-e**). Combining with our optimal copy number model, we modeled the uneven inheritance of ecDNA and even inheritance of HSR to predict changes in copy number distributions over time across populations initialized with different copy numbers. EcDNA(+) cells from distinct copy number groups were predicted to converge at a middle copy number by Day 14, while convergence was not observed in HSR(+) cells at the same timepoint (**Extended Data Fig. 7a-c**). Based on these simulations, we hypothesized that asymmetric ecDNA inheritance, coupled with a non-monotonic (quadratic) relationship between ecDNA copy number and proliferative fitness, is sufficient to drive population-level shifts in ecDNA copy number and convergence toward an optimal range during cancer cell growth. To validate the results of our simulations using an experimental approach, we used FACS to split GBM39-EC and GBM39-HSR cells into low, middle, and high EGFR groups, measured proliferation among cells seeded at uniform initial density across all sorted groups for three consecutive days post-sort, and performed *EGFR* DNA FISH on fixed cells on days 0, 3 and 7 from each group to measure copy number. Since *EGFR* copy number correlates well with EGFR protein expression (**Fig. 1d**), the cell surface EGFR protein abundance is a reasonable estimate of *EGFR* copy number. Using DNA FISH to measure the copy number distributions of the high, middle, and low populations, indeed the copy number distributions among ecDNA(+) cells from different initial copy number groups converged significantly by day 3 and were maintained on Day 7 (**Fig. 3a-b**). Consistent with live-cell imaging data showing that a middle copy number range is associated with the shortest time to division in COLO 320DM cells harboring *MYC* ecDNA (**Fig. 2h**), cells collected from the middle 15% of the initial GBM39-EC and HSR populations also exhibited the best proliferation within the first three days after sorting (**Fig. 3c**). These data warrant that *EGFR* ecDNA copy number is associated with fitness, namely the capacity of ecDNA cancer cells to proliferate, and there is a selection pressure for cells with a middle copy number status at baseline conditions. Notably, convergence in ecDNA(+) cells occurs very rapidly, faster than the time predicted by our simulation (**Extended Data Fig. 7a-b**). This effect may be accounted for by unmodeled cell arrest and death at extreme copy numbers, and incomplete population purity following FACS, together accelerating selection for cells with optimal copy numbers.

**Fig. 3.**
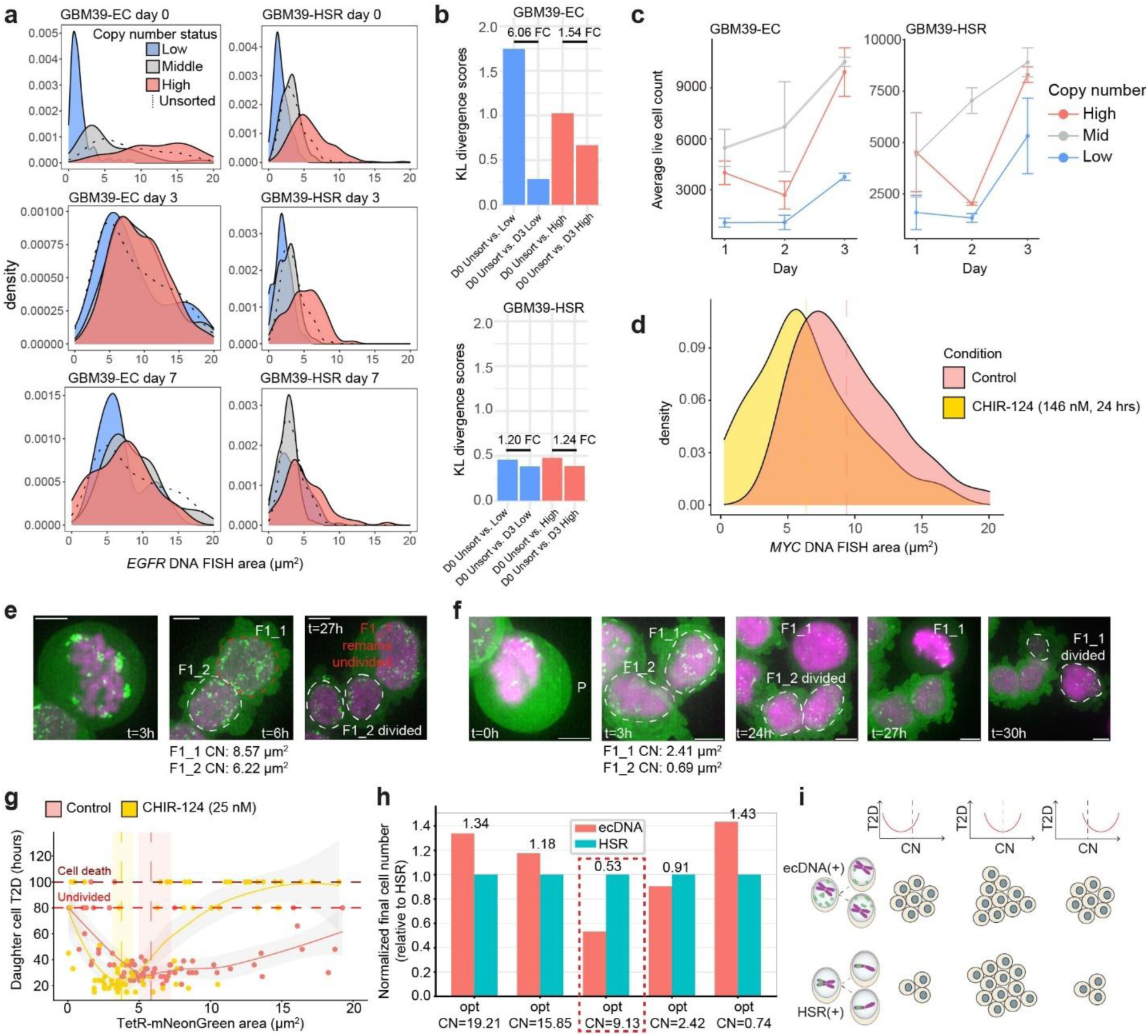
Dynamic optimal ecDNA copy numbers drive cell adaptation. **(a)** Density curves depicting copy number distributions in low, middle, high, and unsorted GBM39-EC and GBM39-HSR cells from *EGFR* DNA FISH on cells at days 0, 3, and 7 post-FACS; n > 30 cells for each group at each timepoint. **(b)** Kullback-Leibler divergence scores between unsorted and low and unsorted and high copy number populations in GBM39-EC and GBM39-HSR at days 0 and 3 post-FACS. **(c)** Average live cells count in low, middle, and high copy number GBM39-EC and GBM39-HSR populations in the first three days post-FACS. Uniform initial seeding density of 3000 cells across all sorted groups. **(d)** Density curves depicting copy number (measured by *MYC* DNA FISH foci area) distributions in untreated- (n=118) versus CHK1-inhibitor-treated (n=46) ecDNA(+) daughter cells (identified by Aurora B Kinase IF). Dashed lines indicate mean copy number of daughter cell population. **(e)** Representative images depicting daughter cells with differential ecDNA copy numbers and cell fates under CHK1-inhibitor treatment. P = parent, F1_x = daughter cells (Generation 1). Scale bars, 5 µm. CN = copy number, measured by TetR-mNeonGreen foci area. Recently divided daughter cells depicted by dashed lines (red indicating poor cell fate). **(f)** Representative images depicting daughter cells with low ecDNA copy numbers successfully undergoing cell division under CHK1-inhibitor treatment. **(g)** Relationship between *MYC* ecDNA copy number (measured by TetR-mNeonGreen foci area) and T2D in TG19 ec under no treatment (n=76 cells) or 25 nM CHK1-inhibitor treatment (n=74 cells) with LOESS-smoothed curves shown in solid lines. Vertical dashed lines indicate the median copy number at which T2D is minimal, derived from 1000 bootstrap resamples, with shaded regions indicating the 95% confidence interval. **(h)** Simulation of ecDNA(+) versus HSR(+) cancer cell growth over ten days in environments requiring different optimal copy numbers and assuming the initial copy number of the single progenitor cell in each group starts with a copy number of 9.13 µm^2^. Red dashed box marks the optimal copy number of cultured TG19 ec cells empirically derived from live-cell imaging. **(i)** Schematic depicting predictions of ecDNA(+) and HSR(+) cell population growth in environments with different optimal copy numbers (minima of red curve). Dashed line in T2D vs. CN plots indicate the initial copy number of the single cell progenitor.

Because ecDNA copy number is linked to opposing fitness pressures, including oncogenic signaling and replication stress-associated DNA damage (**Fig. 1d-g**), we reasoned that optimal copy number is determined by the balance between promoting oncogenic signaling and minimizing DNA damage resulting from replication stress^18^, and that it could change in response to external perturbations, including drug treatment. To assess the impact of replication stress as a regulator of ecDNA abundance, we treated our cells with a known replication stress enhancer, a CHK1 inhibitor. We hypothesized that if replication stress serves as a critical limiting factor for ecDNA abundance, then enhancing replication stress would result in a consistent decrease in ecDNA copy number. Indeed, pharmacological inhibition of CHK1 with CHIR-124^32^ (146 nM = IC50 in COLO 320DM) caused a significant decrease in *MYC* ecDNA copy number within just 24 hours in TG19 ec daughter cells (**Fig. 3d**). While these data revealed rapid population-level copy number reduction, we could not distinguish whether this shift arose from selective loss of high ecDNA copy cells, altered division dynamics, or emergent copy number reduction within surviving lineages. To directly resolve the cellular trajectories underlying this response, we next treated TG19 ec cells with a low dose of the CHK1 inhibitor CHIR-124 (25 nM). This sublethal dose minimized widespread cell death while enabling longitudinal tracking of parental cells and their progeny, capturing dividing, undivided, and dying cells and allowing direct measurement of ecDNA copy number dynamics during drug response. Under this condition, we directly observed the rapid emergence and selection of cells with reduced ecDNA abundance. Notably, cells with low ecDNA copy numbers were more likely to undergo successful and faster division compared to cells with high ecDNA copy numbers that frequently remained undivided or underwent cell death (**Fig. 3e**). By virtue of asymmetric inheritance of ecDNA to daughter cells, cells with low ecDNA abundance and associated increased fitness in the setting of CHK1-inhibitor treatment were also generated de novo, arising from parent cells with higher ecDNA copy numbers (**Fig. 3e-f**). Plotting fitness as a function of ecDNA copy number confirmed a shift in optimal copy number to a significantly lower level compared to that of baseline culture conditions. The median copy number at minimum T2D was −1.98 units lower in CHK1i-treated cells compared to untreated cells, with a 95% bootstrap confidence interval of [-3.71, −1.02]. (**Fig. 3g**). These data demonstrate that replication stress acts as a dominant selective pressure shaping ecDNA copy number dynamics by rapidly eliminating cells with excessive ecDNA burden. This process enriches for pre-existing low copy number ecDNA cells that are resistant to CHK1 inhibition, explaining the results of our previous observation in which surviving cells from SNU16 gastric cancer tumours treated with CHK1i showed remarkably lower *FGFR2* ecDNA copy numbers^18^.

Under an optimal copy number model, simulating cancer cell growth for ten days from a single ecDNA or HSR progenitor with a starting copy number index of 9.13 (**Fig. 2h**) (representing optimal copy number for unchanging, baseline culture conditions) predicted that HSR(+) cells would outcompete ecDNA(+) cells when HSR(+) cells uniformly possess the optimal amplicon copy number. In contrast, when the optimal copy number deviated from this initial value of 9.13, the model predicted that ecDNA(+) cells would outgrow HSR(+) cells, derived from the ability of ecDNA(+) cells to rapidly shift copy numbers via uneven ecDNA inheritance (**Fig. 3h-i**). This framework explains our prior observations that ecDNA(+) cells exhibit superior adaptive capacity relative to HSR(+) cells under selective pressures such as drug treatment and nutrient deprivation^14^. Given the increased complexity and selective pressures of the tumour microenvironment relative to in vitro culture, we next sought to explore how the optimal copy number model accounts for ecDNA copy number dynamics during tumour evolution.

### ecDNA(+) cells exhibit increased tumourigenicity in vivo due to copy number shift

Motivated by prior work demonstrating that ecDNA(+) tumours are associated with worse clinical outcome compared to tumours driven by linear amplification^3,33,1^, we assessed the proliferation of COLO 320DM and COLO 320HSR cancer cells in an in vivo model. We performed two sets of competition assays: one in vitro and another in vivo (**Fig. 4a**). We mixed mCherry-labeled COLO 320DM cells and GFP-labeled COLO 320HSR cells at varying initial ratios (25% ec: 75% HSR, 50% ec: 50% HSR, and 75% ec: 25% HSR) and used flow cytometry to quantify the proportion of DM and HSR cells in each group at 48 hours after initial seeding for in vitro conditions and at 28 days after cells were subcutaneously injected into mouse flanks for in vivo conditions (**Extended Data Fig. 8**). In culture, COLO 320HSR cells outcompeted COLO 320DM cells in every group. In the 50% DM: 50% HSR in vitro group, 70.3% of the total cells were COLO 320HSR (**Fig. 4b**). However, in every mixed DM and HSR group in vivo, COLO 320DM cells outcompeted COLO 320HSR cells. In the 50% DM: 50% HSR in vivo group, 93.1% of the total cells were COLO 320DM (**Fig. 4b**). These findings are consistent with our simulations of ecDNA(+) and HSR(+) cancer cell growth under an optimal copy number model. Specifically, they suggest that HSR(+) cells may harbor an amplicon copy number that is optimal for in vitro culture conditions but suboptimal for proliferation in vivo (**Figure 3i**).

**Fig. 4.**
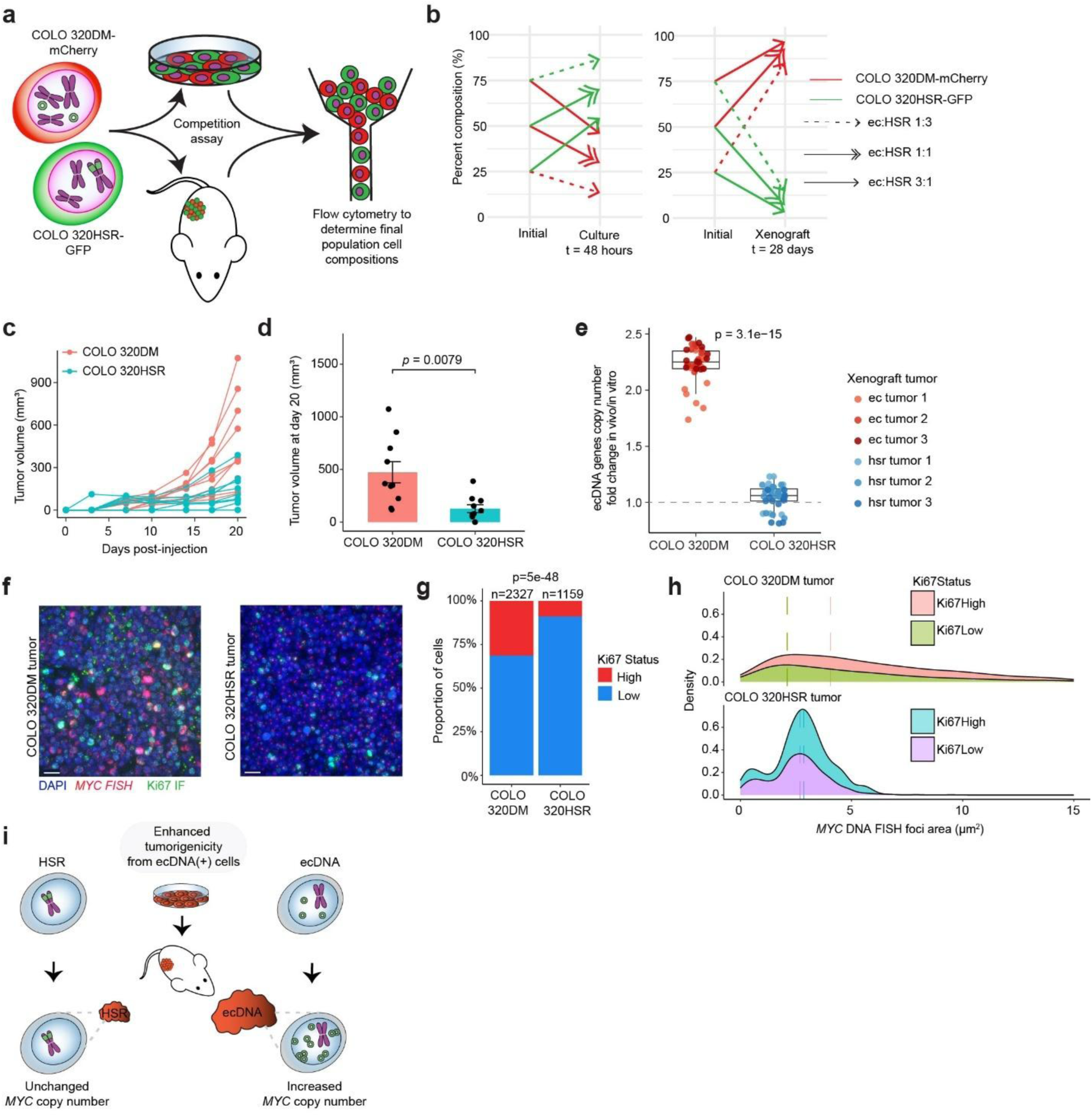
Shift to higher copy number accounts for increased tumourigenicity of ecDNA(+) cells in vivo. **(a)** Schematic illustrating competition assay workflow with COLO 320DM-mCherry and COLO 320HSR-GFP cells. **(b)** Change in cell composition, expressed as the percentage of COLO 320DM-mCherry and COLO 320HSR-GFP cells, at the start and end of competition assays performed in culture and in xenograft mouse models. **(c)** Tumourigenicity plots depicting rate of tumour growth in COLO 320DM (n=5 mice) and COLO 320HSR (n=5 mice) xenograft mouse models. Initial seeding density of 125,000 cells. **(d)** Mean tumour volume on day 20 in COLO 320DM (n=5 mice) and COLO 320HSR (n=5 mice) xenograft mouse models. Statistical significance was assessed using a two-sample independent t-test. **(e)** Copy number fold change of genes encoded on ecDNA between xenograft tumour cells and cultured cells in COLO 320DM and COLO 320HSR. Gene copy numbers were estimated from whole genome sequencing data. Statistical significance was assessed using a two-sided Wilcoxon rank-sum test. **(f)** Representative images from multiplexed FISH for *MYC* and IF for Ki67 on FFPE sections of COLO 320DM and COLO 320HSR xenograft mouse tissue. Initial seeding density of 1000 cells. Scale bars, 20 µm. **(g)** Proportion of COLO 320DM and COLO 320HSR cells by Ki67 expression. Ki67 status was classified as high if the area of Ki67 signal per cell was above 3.079 µm^2^ (2 median absolute deviations) (**Extended Data Fig. 9b**). Difference in the proportion of highly Ki67-expressing cells between samples was assessed using a chi-squared test. **(h)** Density curves depicting proportion of cells vs. *MYC* copy number (measured by *MYC* DNA FISH area) for COLO 320DM (n=2327) and COLO 320HSR (n=1159) tissue cells, stratified by Ki67 expression status. Dashed lines indicate the mean copy number for each group. **(i)** Schematic summarizing that the increased tumourigenicity of COLO 320DM cells in vivo can be accounted for by their ability to shift to a higher optimal *MYC* copy number.

In vivo tumourigenicity assays revealed that COLO 320DM tumours exhibited increased rate of tumour growth compared to COLO 320HSR tumours (**Fig. 4c**). The mean tumour volume on day 20 was significantly higher in COLO 320DM (473.3 mm^3^) compared to COLO 320HSR (127.6 mm^3^) mouse xenograft models (**Fig. 4d**). Copy numbers (measured from whole genome sequencing) of genes encoded on ecDNA were significantly increased in COLO 320DM mouse xenograft tumour tissue (average fold change = 2.23) compared to COLO 320HSR mouse xenograft tumour tissue (average fold change = 1.05) (**Fig. 4e**). Metaphase DNA FISH for *MYC* on dissociated tumour cells showed no evidence of integration of ecDNA as HSRs in COLO 320DM mouse tumours nor amplicon switching from HSR to ecDNA in COLO 320HSR mouse tumours (**Extended Data Fig. 9a**).

Based on these results, we hypothesized that COLO 320HSR cells whose *MYC* copy numbers are uniform throughout the population possess their optimal copy number for culture conditions, but the optimal copy number of *MYC* for tumourigenesis increases in vivo. We reasoned that COLO 320HSR cells were less tumourigenic in vivo since they cannot readily change their *MYC* copy number, while the uneven inheritance of ecDNA enabled the COLO 320DM cell population to shift to a higher copy number associated with better proliferation in vivo. Combined FISH for *MYC* and IF for Ki67 in FFPE sections of COLO 320DM and COLO 320HSR mouse xenograft tumour tissue revealed a greater proportion of cycling cells in COLO 320DM compared to COLO 320HSR mouse xenograft tumour tissue, consistent with tumourigenicity results (**Fig. 4f-g**). Copy number associated with the greatest number of cycling cells in COLO 320DM mouse xenograft tumour tissue was higher than the copy number associated with the greatest number of cycling cells in COLO 320HSR mouse xenograft tumour tissue (**Fig. 4h, Extended Data Fig. 9b**). The mean copy number was not significantly different between COLO 320HSR Ki67-high versus Ki67-low cells, consistent with the idea that each COLO 320HSR daughter cell inherits equivalent *MYC* copy number during cell division, maintaining copy number homogeneity. Collectively, these results show that the increased tumourigenicity of COLO 320DM cells can be accounted for by the ability to shift to a higher optimal *MYC* copy number associated with increased proliferation in vivo.

## DISCUSSION

Recent advancements in sequencing technologies have shed light on the widespread prevalence of ecDNA in cancer^1,2^ and its contribution to escalating malignant potential via increased intratumoural heterogeneity^14,34^. Despite this progress, the specific impact of ecDNA copy number on shaping cancer cellular phenotypes remains a central and unresolved question. Here, using single-cell-resolved approaches, we identify key determinants of ecDNA copy number dynamics. Single-cell multiomics analyses show that ecDNA abundance is constrained by opposing selection pressures, including enhanced oncogenic signaling and elevated cellular stress, which are conserved across diverse ecDNA(+) cancers. Transgenerational live-cell imaging demonstrates that asymmetric inheritance of ecDNA directly generates heterogeneity in copy number and proliferative capacity. Moreover, we show that cancer cells dynamically shift ecDNA copy number toward new optimal ranges in response to environmental change, thereby enhancing tumourigenicity and driving tumour evolution.

A key strength of this study is the application of live-cell imaging to directly interrogate ecDNA dynamics in living cells. By establishing a near-isogenic pair of fluorescently labeled ecDNA and chromosomal amplicon (HSR) cell lines, we were able to track ecDNA copy number and time to division across multiple generations. Long-term live-cell imaging provides insight into the kinetics at which ecDNA-conferred copy number heterogeneity develops, demonstrating that a single ecDNA(+) progenitor can give rise to a very diverse population of cells within few cell divisions, accelerating subclone development. We have shown that there is immense heterogeneity of growth patterns within a single population of cancer cells, particularly those with ecDNA. Notably, the isogenic pair of live-cell imaging cell lines highlights functional cell biology that is unique to ecDNA(+) cancer cells, including increased variation in the ability to divide and time to next division. These observations motivate future studies that combine ecDNA reporters with additional functional sensors, such as DNA damage or replication stress reporters, to directly assess the causal impact of damaged ecDNA in shaping cancer cell fitness.

Our findings further demonstrated that ecDNA(+) cancer cell fitness is strongly coupled to copy number, and optimal copy number is a common feature of several ecDNA(+) cell lines, including COLO 320DM, PC3 DM, and GBM39-EC. Although the precise numerical optima differ across measurement modalities, we consistently observe a middle optimal range associated with faster division and increased proliferation at baseline across single-cell multiomics analyses, live-cell imaging, protein-based sorting, and multiplexed IF-FISH. We find that elevated ecDNA copy number amplifications are associated with a trade-off, as DNA damage signaling may limit the extent to which ecDNA-driven cancer cells can enhance oncogenic signaling. For instance, a cancer cell carrying a high copy number of ecDNA also increases the probability that one of the ecDNA molecules undergoes stochastic DNA damage from replication stress, leading to arrest or cell death; hence the reward to risk ratio for harboring ecDNA changes with each additional copy. By combining single-cell multiomics sequencing and multiplexed imaging across multiple cancer cell lines and a human patient glioblastoma sample, we have shown that critical determinants of optimal copy number include oncogenic pathways and replication stress-associated DNA damage, which curbs ecDNA abundance from continuously increasing without limit. Additionally, our study makes increasingly clear that copy number shifts are critical for ecDNA(+) cancer cell adaptation to different environments with different optimal copy number ranges.

This framework provides a mechanistic explanation for several prior associations between ecDNA copy number shifts and therapeutic resistance, revealing that intrinsic ecDNA copy number heterogeneity gives rise to subpopulations with specific copy numbers that are resistant to treatment^35^. In glioblastoma, ecDNA(+) GBM39-EC cells tolerate erlotinib more effectively than cells with chromosomally integrated *EGFR* amplifications, in part because the surviving ecDNA(+) population shifts to a lower *EGFR* copy number under treatment pressure^14^. Other recent work similarly revealed that *MYCN* copy number decreases upon treatment with genotoxic stressors in neuroblastoma cells^17^. In contrast, SNU16 gastric cancer cells harboring *FGFR2* and *MYC* on distinct ecDNAs increase *FGFR2* copy number upon infigratinib exposure^18^. These observations suggest that ecDNA copy number modulation is drug-specific and depends on how a specific treatment perturbs the balance between oncogenic signaling and cellular stress. Our findings therefore support that a combinatorial approach with therapies that synergistically target diverse copy number ranges, such as coupling a replication stress enhancer like a CHK1-inhibitor with an oncogene suppressor like infigratinib in SNU16 cells^18^, as an effective strategy against ecDNA(+) cancers.

Other strategies that may hold promise against ecDNA(+) cancers include therapies that can target or silence retention elements and factors critical for the tethering of ecDNA to mitotic chromatin to decrease ecDNA copy number^36^. We believe that the TG19 ec and TG19 HSR isogenic live-cell imaging pair of cell lines will serve enormous utility for identifying ecDNA copy number-dependent sensitivities from high-throughput screens. The fluorescent-based, real-time copy number readout eliminates the need for performing additional experimentation to measure ecDNA copy number after treatment. Being able to efficiently collect information about changes in ecDNA copy number or tethering in response to varied drug treatments will be essential for determining and tailoring therapeutic combinations that will combat resistance arising from shifts in ecDNA abundance.

Beyond drug response, ecDNA copy number plasticity also contributes to tumour progression in vivo. We found that ecDNA(+) cells exhibit greater tumourigenicity in vivo than HSR(+) cells, and this increased tumourigenicity strongly correlates with an increase in ecDNA copy number to a new optimal level. It is unclear why the in vivo tumour microenvironment selects for increased copy number. Given that *MYC* detected by single-cell RNA sequencing shows high expression in COLO 320DM xenograft cells but its expression is massively decreased in COLO320 HSR xenograft cells^37^, our hypothesis is that MYC was downregulated in vivo and subsequently, cancer cells with ecDNA compensate for decreased *MYC* expression by increasing their copy number. The mechanistic explanations behind inhibitory *MYC* feedback in vivo warrant further study. Other work has shown how copy number influences subclonal expansion and metastatic potential in lung cancer^41^. These findings suggest that metastatic niches may require a different optimal copy than the primary tumour site, and ecDNA(+) cells can more effectively seed at these secondary sites by shifting their copy number. Future studies to track ecDNA levels with varying cargoes over space and time, including at metastatic sites in treated patients will be critical for determining how the dynamic nature of ecDNA contributes to metastases. While we establish oncogenic copy number shifts as a major contributor to ecDNA(+) cell adaptation, other factors, including previously reported associations between ecDNA and lower immunogenicity^1,42–45^ or prevalent ecDNA-borne oncogene fusions^37^, may account for increased tumourigenicity of ecDNA(+) cells in vivo. Oncogene cargo itself could also potentially play a role, including in which oncogenes and/or immunomodulatory genes and regulatory elements are amplified. Future studies will be needed to elucidate how immunomodulatory^1^ and regulatory^5,46^ ecDNA molecules play unique roles in the tumour microenvironment to potentially contribute to tumour progression and metastasis.

In summary, our results establish ecDNA copy number plasticity as a fundamental mechanism by which cancer cells generate phenotypic heterogeneity, adapt to therapeutic and microenvironmental pressures, and evolve toward increased malignancy. These findings underscore the importance of early ecDNA detection in patients and highlight dynamic ecDNA copy number regulation as a promising vulnerability for the development of next-generation combination therapies.

## Acknowledgements

We thank the members of the Mischel and Chang labs for discussion. This project was supported by Cancer Grand Challenges CGCSDF-2021\100007 with support from Cancer Research UK and the National Cancer Institute (P.S.M., H.Y.C.). A.G. was supported by Stanford University Medical Scientist Training Program grant T32-GM145402. X.Y. is a Damon Runyon Fellow supported by the Damon Runyon Cancer Research Foundation (DRG-2474-22). Work on ecDNAs in the Ventura lab is supported by grants from NIH-NCI (P30 CA008748 to the MSK Cancer Center, and R01 CA282913 to AV), the Cancer Grand Challenge Initiative, the American Cancer Society Discovery Boost grant, and The Mark Foundation for Cancer Research ASPIRE grant. B.W. is supported by a UKRI Future Leaders Fellowship (grant no. MR/V02342X/1). H.Y.C. was an Investigator of the Howard Hughes Medical Institute.

## Author Contributions

A.G., S.Z., E.J.C., H.Y.C., and P.S.M. designed the study. R.L. prepared single-cell multiomics sequencing libraries. S.Z. analyzed single-cell multiomics data. A.G. and J.T. performed fixed interphase cell IF-DNA FISH staining and image acquisition and analysis. A.G., E.J.C., and X.Y. prepared reagents for and performed live-cell imaging. A.G. and E.J.C. processed and analyzed live-cell imaging data. S.Z. and V.S. performed flow cytometry. S.Z. and L.Y. performed copy number-dependent modeling of cancer cell growth. E.J.C., S.Z., and I.T.L.W. generated, maintained, and monitored xenograft mouse models. A.G. performed metaphase DNA FISH. A.G. and M.Y. performed IF-DNA FISH on FFPE tissue. A.V., B.W., H.Y.C, and P.S.M. guided data analysis and provided feedback on experimental design. A.G., S.Z., H.Y.C., and P.S.M. wrote the manuscript with input from all authors. H.Y.C. and P.S.M. supervised the project.

---

**Extended Data Fig. 1.**
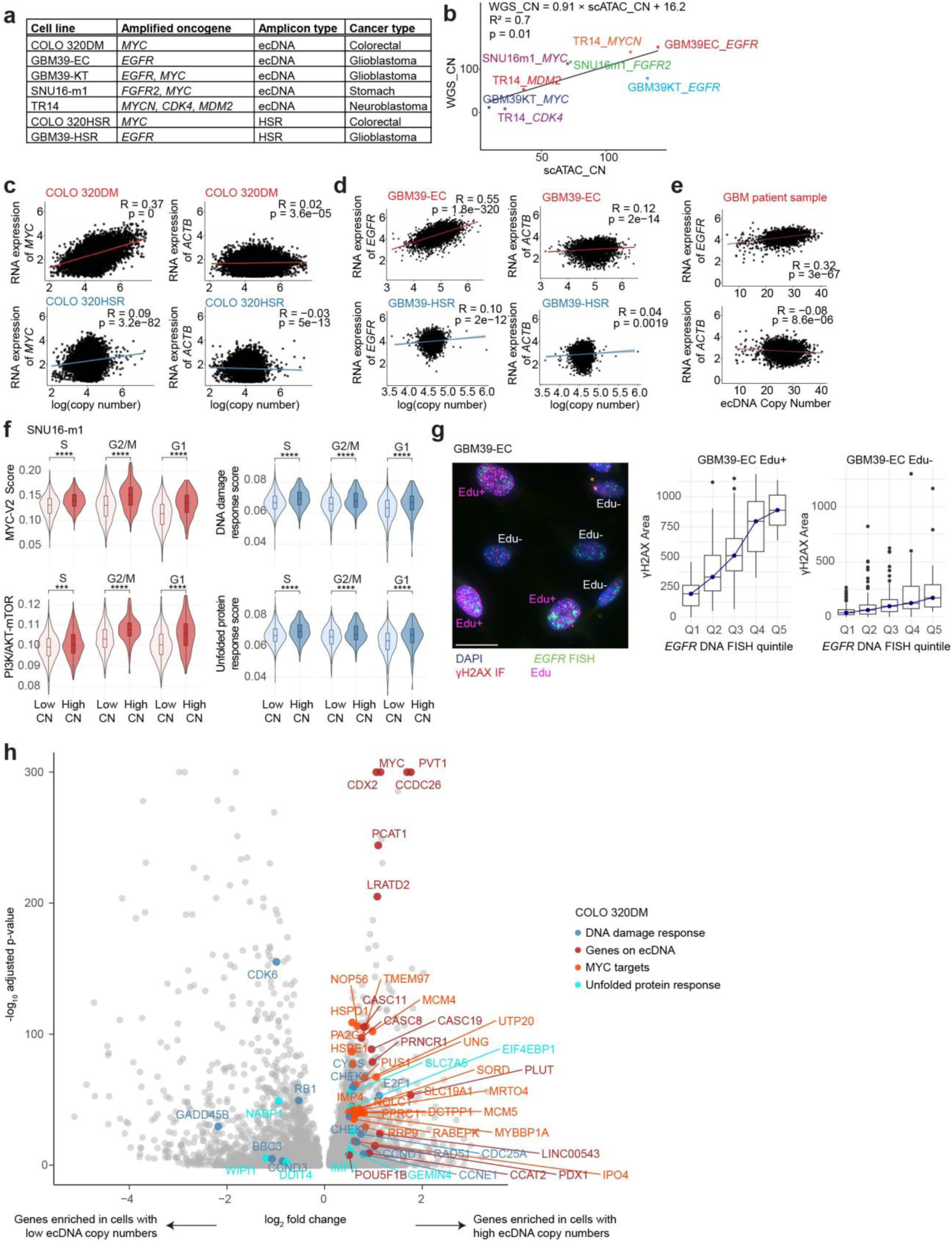
Copy number and transcript heterogeneity are generalizable across ecDNA(+) samples. **(a)** Table of cell lines used for single-cell multiomics analyses. **(b)** Pearson correlation scatterplot demonstrating concordance between ecDNA oncogene copy number estimated by whole genome sequencing and by scATAC-seq. Colors used to match point to text label. **(c)** Pearson correlation scatterplots between copy number (measured by scATAC-seq) and transcript expression (measured by scRNA-seq) for *MYC* and housekeeping gene *ACTB* in COLO 320DM and COLO 320HSR. **(d)** Pearson correlation scatterplots between copy number and transcript expression for *EGFR* and *ACTB* in GBM39-EC and GBM39-HSR. **(e)** Pearson correlation scatterplots between copy number and transcript expression for *EGFR* and *ACTB* in a human patient glioblastoma sample. **(f)** Distributions of pathway expression scores for MYC-V2 targets, PI3K_AKT_MTOR expression, DNA damage response, and unfolded protein response between cells with high and low ecDNA copy numbers across different cell cycle stages in the SNU16-m1 cell line, which has both *MYC* and *FGFR2* amplified on ecDNA. Statistical significance was assessed using a two-sided Wilcoxon rank-sum test. **(g)** Representative image of combined IF for γH2AX and DNA FISH for *EGFR* in GBM39-EC cells with Edu labeling to distinguish replicating cells (in S-phase) from non-replicating cells. Scale bar, 20 µm (left). Distribution of γH2AX expression (right) across *EGFR* ecDNA copy number quintiles among Edu-positive (n=220) and Edu-negative (n=605) cells. **(h)** Volcano plot demonstrating differentially expressed genes between cells with low and high ecDNA copy numbers in COLO 320DM. Genes found on ecDNA and genes in the MYC-V2 targets, DNA damage response, and unfolded protein response pathway signatures are labeled.

**Extended Data Fig. 2.**
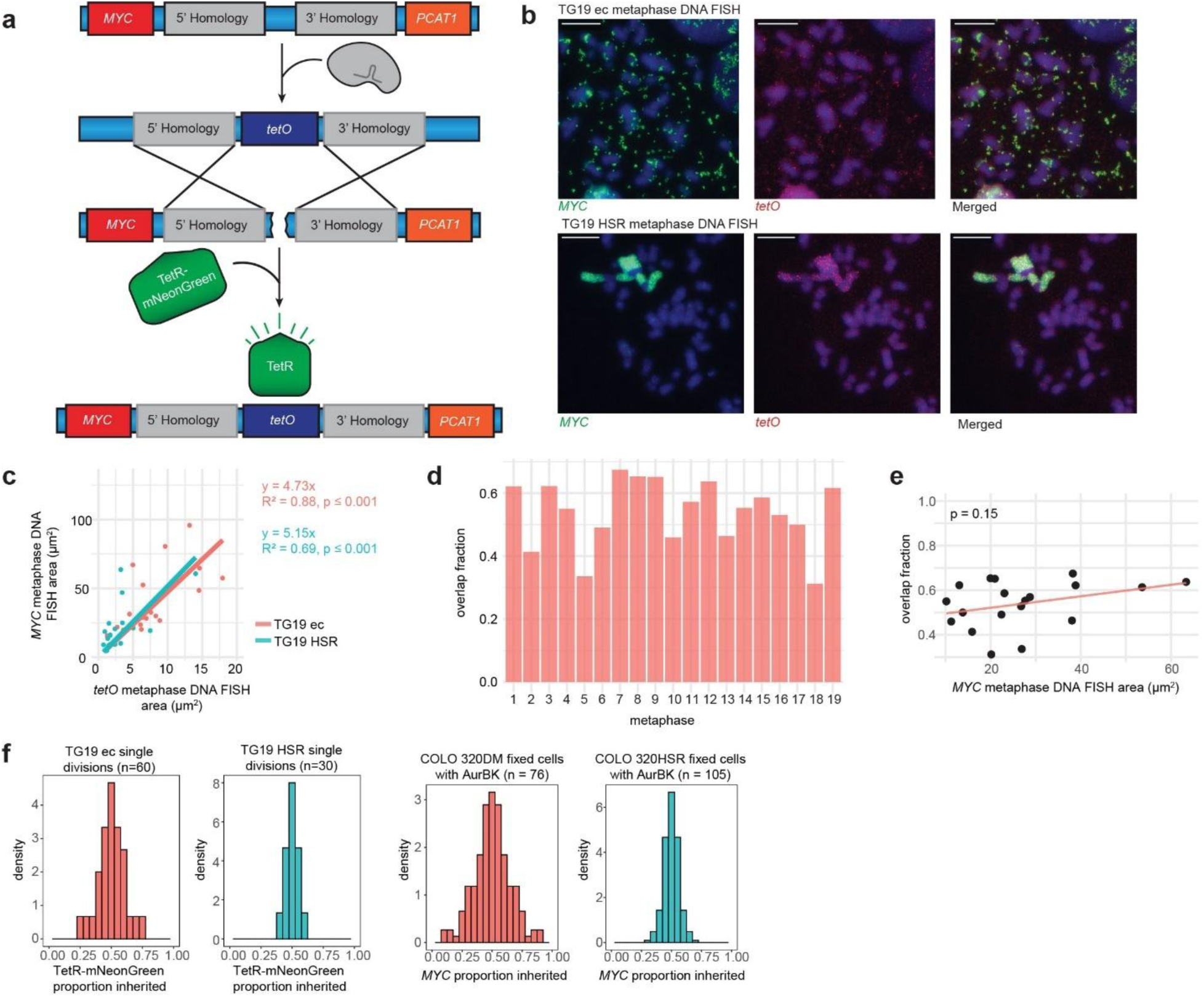
TetR-mNeonGreen foci is a reliable proxy for *MYC* copy number. **(a)** Schematic depicting CRISPR-Cas9-mediated knock-in of a *tetO* array (96x) into the intergenic region between *MYC* and *PCAT1* and subsequent visualization of *tetO* upon binding to TetR-mNeonGreen. We named the engineered COLO 320DM-*tetO* and COLO 320HSR-*tetO* lines TG19 ec and TG19 HSR, respectively. **(b)** Representative images of metaphase DNA FISH (*tetO* in red and *MYC* in green) in TG19 ec and TG19 HSR. Scale bars, 10 µm **(c)** Pearson correlation scatterplot depicting correlation between *tetO* area and *MYC* area measured from TG19 ec (n=19) and TG19 HSR (n=20) metaphase DNA FISH images. **(d)** Fraction of overlap between *tetO* and *MYC* foci for each TG19 ec metaphase DNA FISH image. **(e)** Pearson correlation scatterplot depicting correlation between *MYC* copy number (measured by *MYC* DNA FISH foci area) and overlap fraction. **(f)** Frequency histograms depicting proportion of *MYC* inherited (measured by TetR-mNeonGreen foci area) to daughter cells in dividing TG19 ec cells (n=60 divisions) and TG19 HSR cells (n=30 divisions) captured by live-cell imaging (left). Frequency histograms depicting proportion of *MYC* inherited (measured by *MYC* DNA FISH foci area) to daughter cells (indicated by Aurora B Kinase IF) in fixed COLO 320DM (n=76 divisions) and COLO 320HSR cells (n=105 divisions) (right).

**Extended Data Fig. 3.**
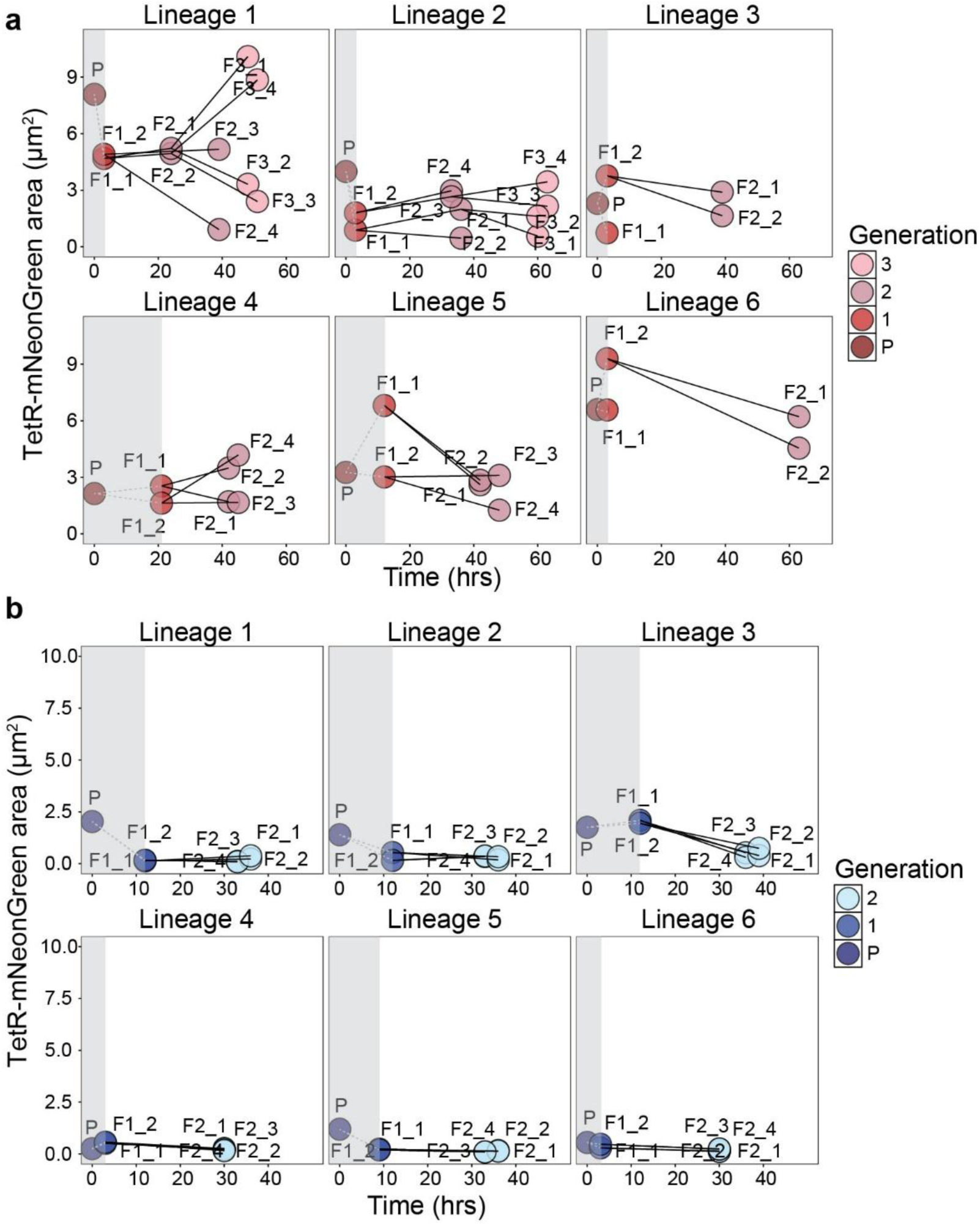
TG19 ec cells show greater copy number and division time heterogeneity than TG19 HSR cells. Lineage plots depicting *MYC* copy number (measured by TetR-mNeonGreen foci area) and T2D for six unique lineages in **(a)** TG19 ec and **(b)** TG19 HSR captured in one live-cell imaging experiment. Live-cell imaging was conducted for 63 hours with a 3-hour timestep. Shaded gray areas denote the time that elapsed before the single progenitor parent cell first divided. At the beginning of image acquisition, parent cells were mitotic. Thus, the time between the parent and F1 cells does not represent the true cycling time from G1 to cell division.

**Extended Data Fig. 4.**
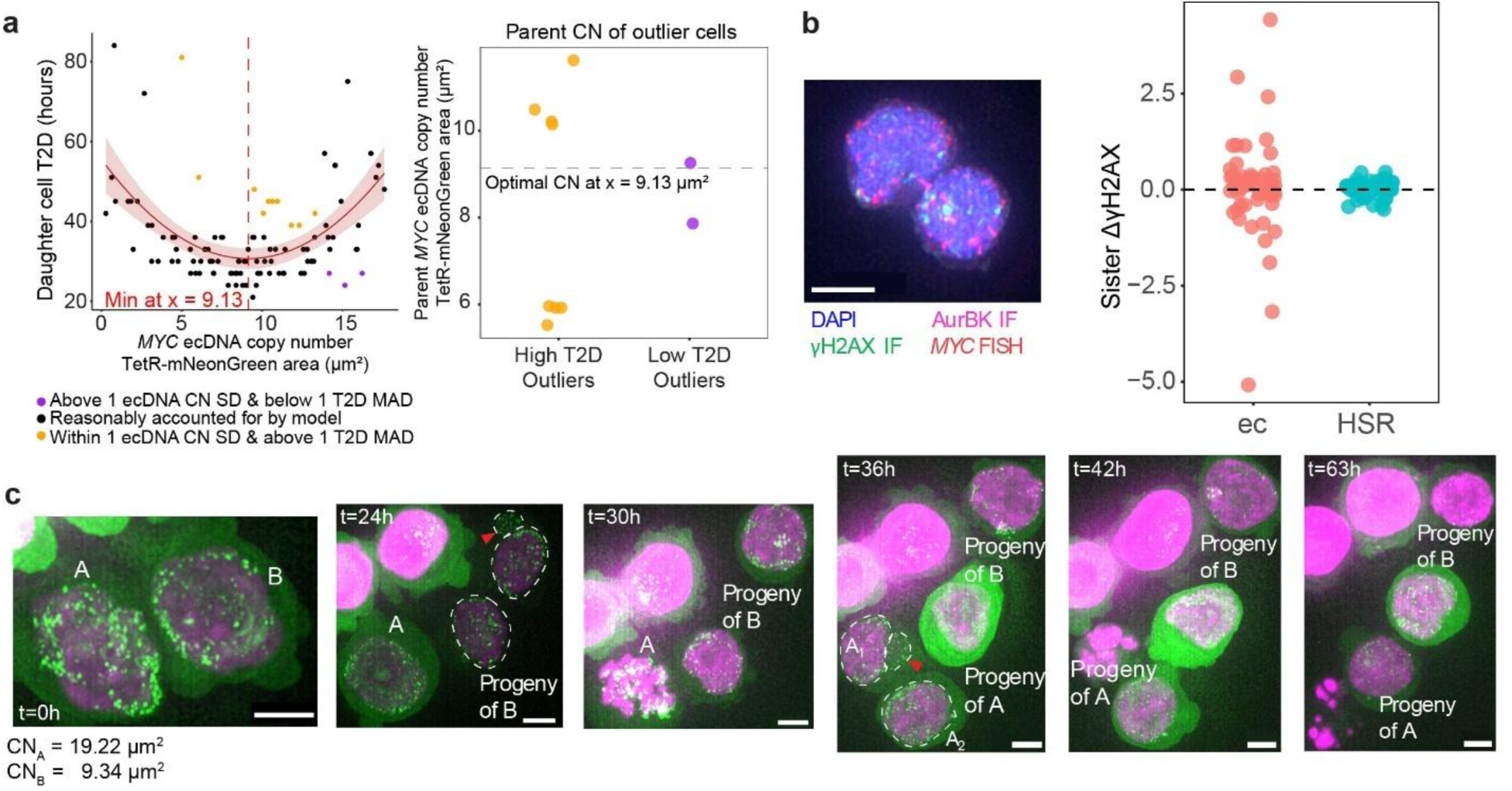
Uneven inheritance of DNA damage impacts daughter cell fitness. **(a)** Relationship between ecDNA copy number (measured by TetR-mNeonGreen foci area; CN = copy number) and T2D in TG19 ec. Unexpectedly high T2D outliers are highlighted in orange, and unexpectedly low T2D outliers are highlighted in purple. High T2D outliers were identified as cells whose TetR-mNeonGreen foci area was within one standard deviation (SD) from the minimum value at TetR-mNeonGreen foci area = 9.13 µm^2^ but whose T2D was above one median absolution deviation (MAD) from the median of 33 hours. Low T2D outliers were identified as cells whose TetR-mNeonGreen foci area was outside of one SD from TetR-mNeonGreen foci area = 9.13 µm^2^ but whose T2D was below 1 MAD from the median of 33 hours (left). By retrospective lineage tracking, the parent cell of each outlier was identified, and the *MYC* ecDNA copy number of the parent was estimated by dividing the measured TetR-mNeonGreen foci area by two to account for genome doubling after S-phase (right). Cells in which ecDNA copy number could not reliably be measured and estimated in the parent cell due to chromatin condensation at time of image acquisition were excluded from analysis. **(b)** Representative image of combined IF for Aurora B Kinase and γH2AX and DNA FISH for *MYC* in TG19 ec. Scale bar, 5 µm (left). Quantification of γH2AX difference between sister cells in TG19 ec (n=59 sister cell pairs) and TG19 HSR (n=63 sister cell pairs). Each dot represents the difference between two sister cells (right). **(c)** Representative images from live-cell imaging of TG19 ec cells with ecDNA sequestered in micronuclei (denoted by red arrows). Copy number (CN) measured by TetR-mNeonGreen foci area. Scale bars, 5 µm.

**Extended Data Fig. 5.**
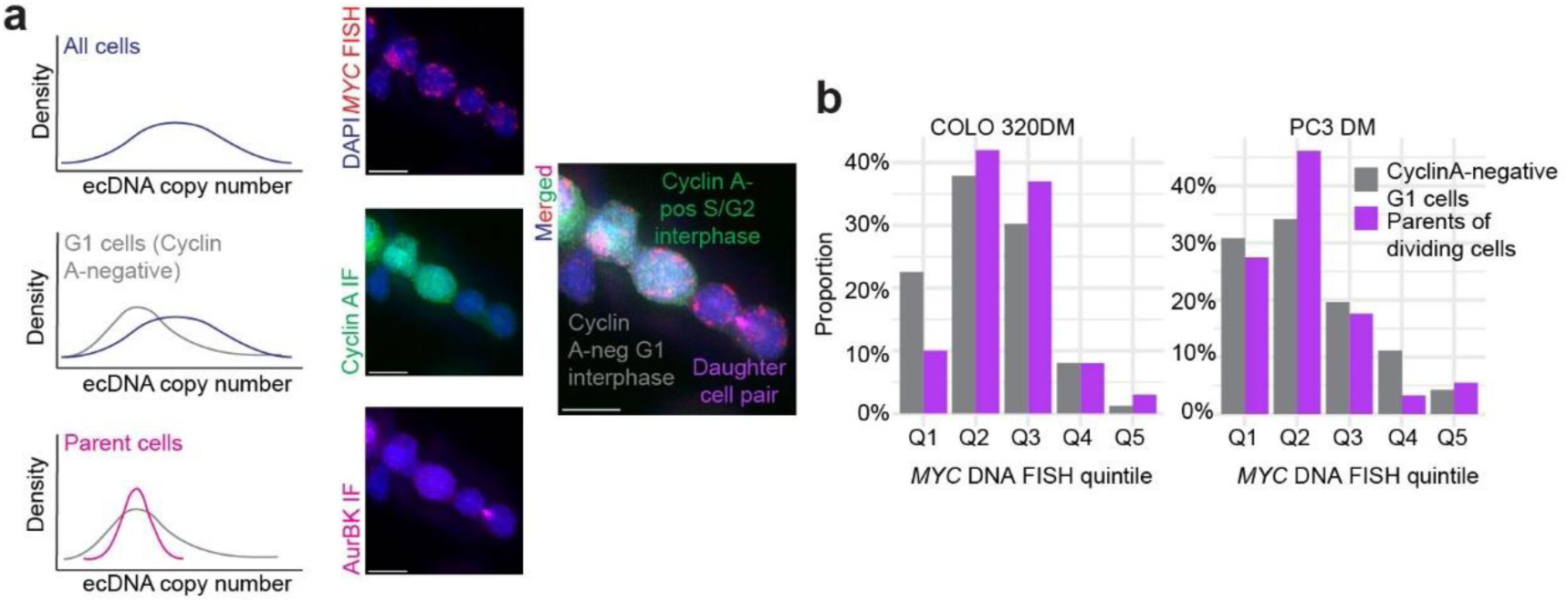
Evidence of optimal copy number across ecDNA(+) cell lines. **(a)** Schematics depicting hypothesized ecDNA copy number distributions of all cells, G1 cells, and parent cells giving rise to daughter cells (left). Representative images from combined IF for Aurora B Kinase and Cyclin A and DNA FISH for *MYC* in COLO 320DM. Cyclin A was used as a marker to identify G1 interphase cells, whose copy number should theoretically be 1n before entering S-phase and undergoing genome doubling. Aurora B Kinase was used as an identification marker for recently divided daughter cells. Copy number of parents of daughter cells was estimated by taking the average of the daughter cells’ *MYC* DNA FISH foci areas. Copy numbers of parent cells were compared against copy numbers of G1 interphase cells to avoid high copy number bias arising from cells past S-phase and with replicated DNA content. Scale bars, 10 µm (right). **(b)** Histograms depicting proportion of G1 interphase and parent cells in each copy number quintile in COLO 320DM and PC3 DM.

**Extended Data Fig. 6.**
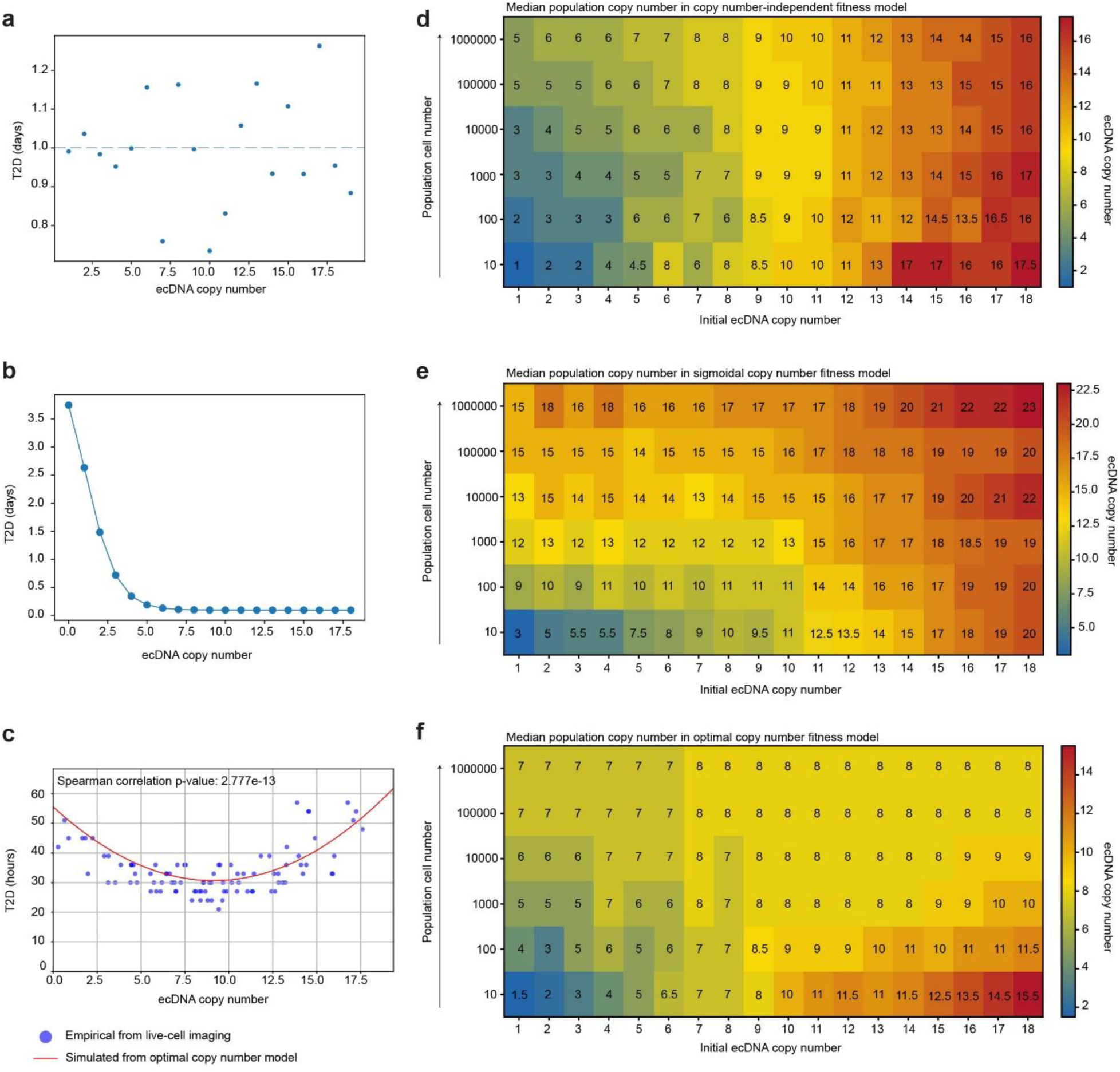
Simulating ecDNA(+) cell population copy number shifts among alternative functional relationships between copy number and T2D. Prediction of T2D vs. ecDNA copy number under a **(a)** horizontal model, which assumes an independent relationship between copy number and fitness, **(b)** sigmoid-plateau model, which assumes T2D decreases and then plateaus as copy number increases, and **(c)** quadratic or optimal copy number model (red curve). **(d)** Median copy number predictions under a **(d)** horizontal model, **(e)** sigmoid-plateau model, and **(f)** optimal copy number model of an ecDNA(+) cell population as it grows from 1 to 1000000 cells. Each column on the x-axis represents the initial copy number of the single cell progenitor.

**Extended Data Fig. 7.**
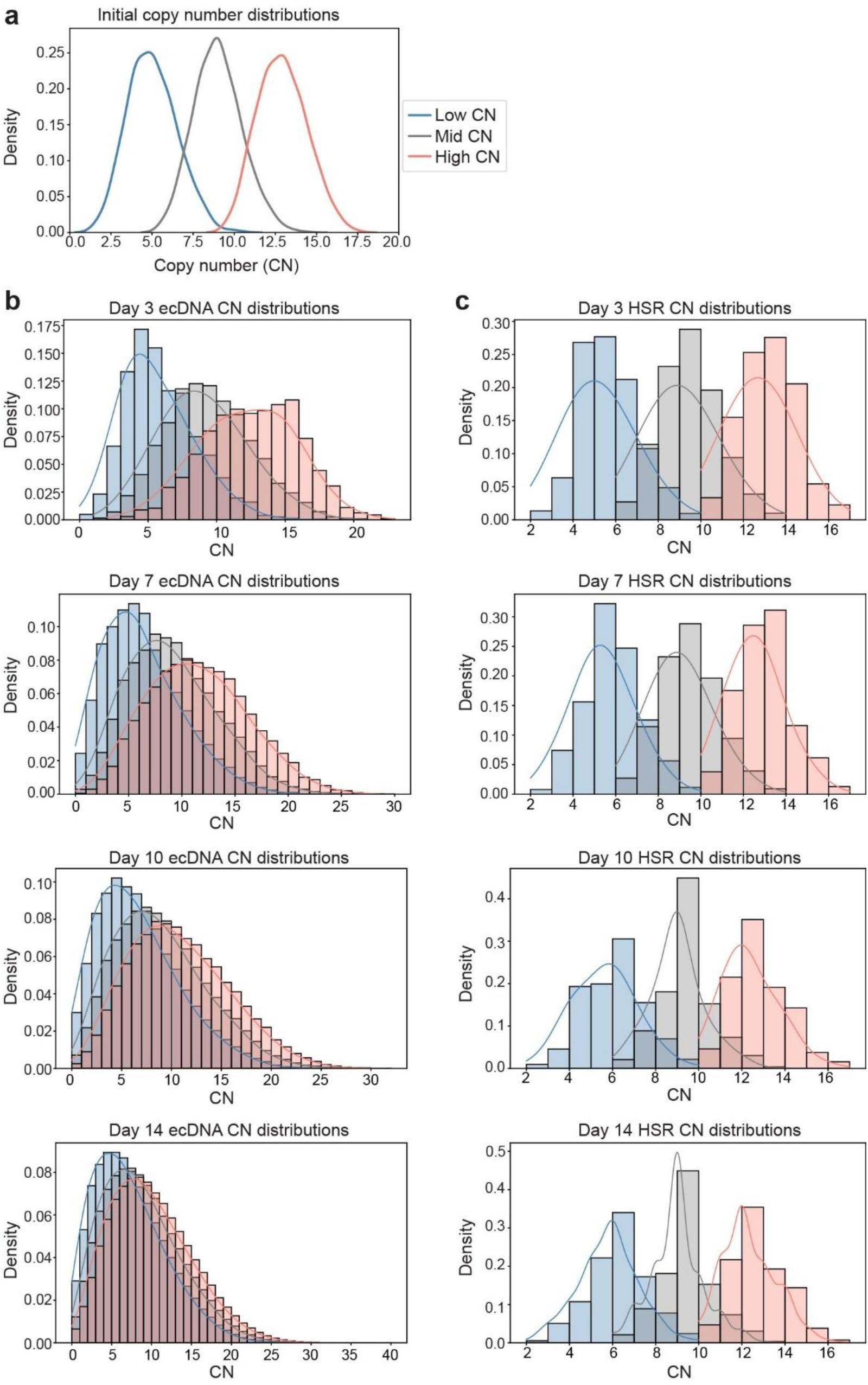
Using optimal copy number model and inheritance patterns to simulate shifts in copy number distributions of ecDNA(+) and HSR(+) cell populations. **(a)** Kernel density estimate plot showing initial copy number (CN) distributions (low, middle, and high) of cancer cells. **(b)** Histograms overlaid with density curves depicting change in copy number distributions of the initially low, middle, and high copy number ecDNA(+) cell populations over time (from Day 3 to Day 14). **(c)** Histograms overlaid with density curves depicting change in copy number distributions of the initially low, middle, and high copy number HSR(+) cell populations over time (from Day 3 to Day 14).

**Extended Data Fig. 8.**
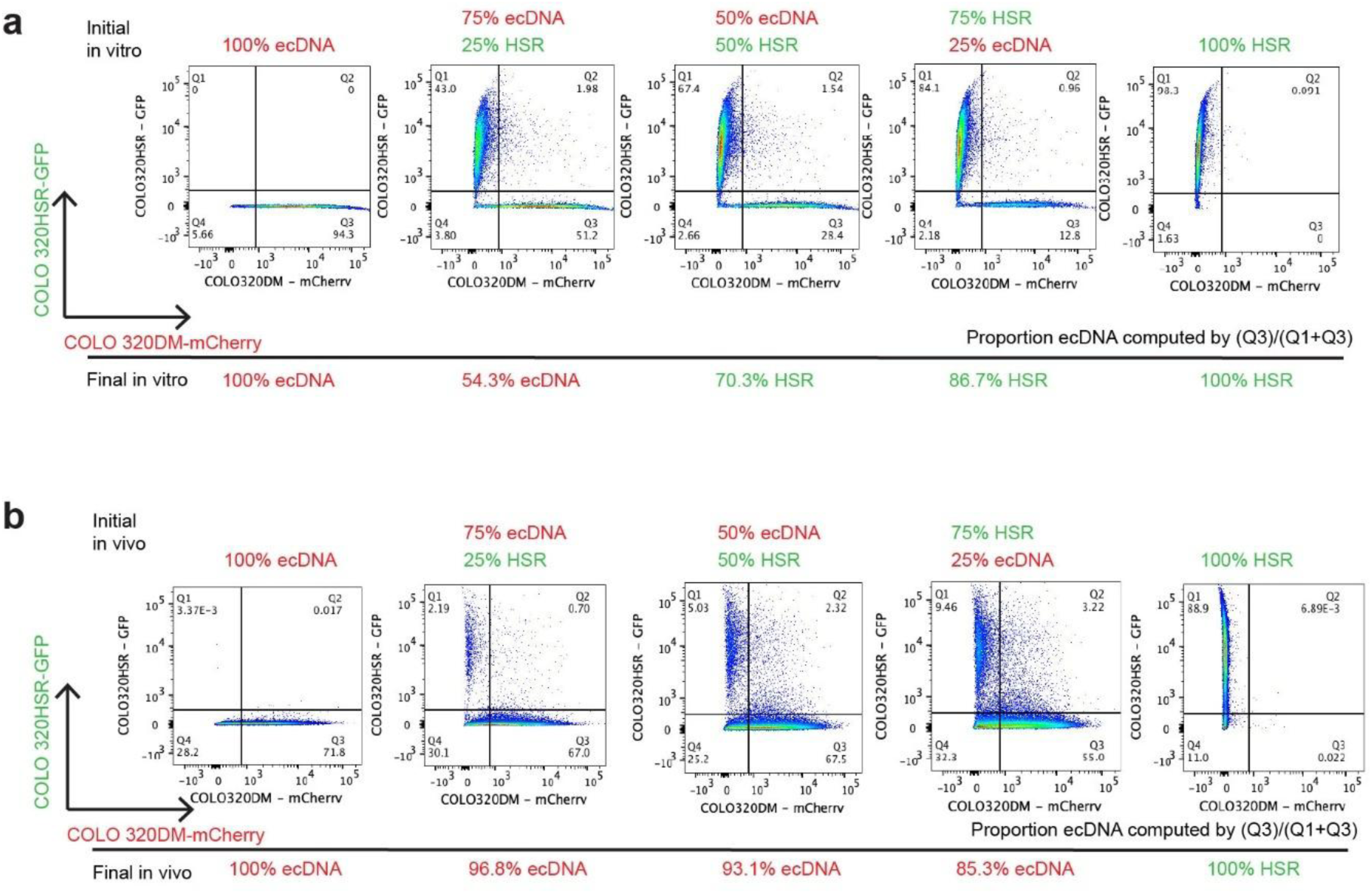
Gating strategy and flow cytometry plots from competition assays. Flow cytometry plots of single and mixed COLO 320DM-mCherry and COLO 320HSR-GFP populations from **(a)** in vitro and **(b)** in vivo competition assays. Gating strategy was established from single populations.

**Extended Data Fig. 9.**
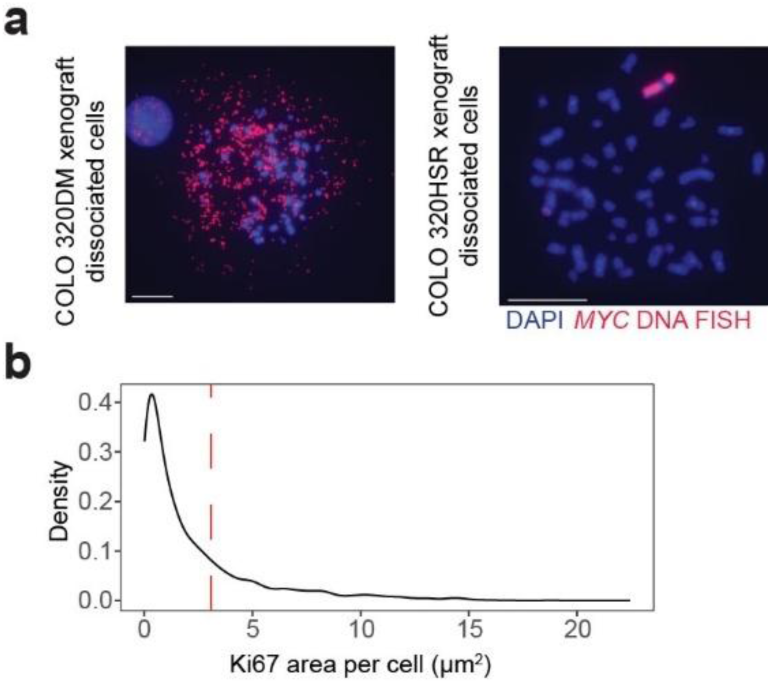
Staining in mouse xenograft cancer cells. **(a)** Metaphase DNA FISH (*MYC*) in dissociated cells from COLO 320DM and COLO 320HSR mouse xenograft tissue. **(b)** Density curve of Ki67 expression from multiplexed IF (Ki67) and FISH (*MYC*) in COLO 320DM and COLO 320HSR FFPE mouse xenograft tissue sections. The dashed red line at 3.079 µm^2^ (2 median absolute deviations) marks the threshold for Ki67-High versus Ki67-Low status.

## METHODS

### Antibodies and reagents

Antibodies were procured as follows: γH2AX (Millipore, catalogue no. 05-636), cyclin A (BD Biosciences, no. 611268), EGFRvIII mAb 806^47^ (see ref), AKT [p Ser 473] (Novus Biologicals, no. AF887), Aurora B kinase (Thermo Fisher Scientific, no. A300-431A), and Ki67 (abcam, no. ab15580). Chemicals were procured as follows: CHIR-124 (Selleckchem, no. S2683).

### Cell culture

COLO 320DM and COLO 320HSR (colorectal cancer) and the parental PC3 (prostate cancer) cell lines were obtained from ATCC. PC3 DM and PC3 HSR lines were isolated by the Mischel lab through single-cell expansions of the parental cell line. COLO 320DM, COLO 320HSR, PC3 DM, PC3 HSR, TG19 ec and TG19 HSR cells were cultured in 4.5 g l^−1^ glucose-formulated Dulbecco’s Modified Eagle’s Medium (Corning) supplemented with 10% fetal bovine serum (FBS; Gibco) and 1% penicillin-streptomycin-glutamine. GBM39 (glioblastoma) cells were patient-derived neurosphere cell lines. GBM39-EC and GBM39-HSR single clones were isolated by Cell Microsystems using CellRaft AIR® System. GBM39-EC and GBM39-HSR cells were cultured in Dulbecco’s Modified Eagle’s Medium/F12 (Gibco, 11320-033) with 1x B-27 supplement (Gibco, 17504044), 1% penicillin–streptomycin, GlutaMAX (Gibco, 35050061), human epidermal growth factor (EGF, 20 ng ml^−1^; Sigma-Aldrich, E9644), human fibroblast growth factor (FGF, 20 ng ml−1; Peprotech, AF-100-18B) and heparin (5 μg ml−1; Sigma-Aldrich, H3149-500KU). All the cells were maintained at 37 °C in a humidified incubator with 5% CO_2_. Cell lines routinely tested negative for mycoplasma contamination.

### IF and DNA FISH staining on cultured cells

Combined IF and DNA FISH were performed on cells grown on coverslips or fixed cells cytospun onto glass slides. Coverslips: Coverslips were coated with 100 µg ml^−1^ poly-l-lysine or 10 µg ml^−1^ laminin or 10 µg ml^−1^ fibronectin for 1 h at 37 °C before seeding cells. Cells were left at 37 °C for approximately 24 hours to adhere to coated coverslips. Where indicated, EdU (Click-iT Plus EdU Alexa Fluor 647 Imaging Kit. Invitrogen, catalogue no. C10640) was added at 10 µg ml^−1^ 30 min before fixation with 4% paraformaldehyde (PFA) for 10 min. Cytospun cells: Cells were treated with trypsin (adherent) or TrypLE (neurosphere) and collected and fixed with 4% paraformaldehyde (PFA) for 10 min. Fixed cells were then diluted to 60 to 100K cells per 300 µL of PBS and cytospun onto glass slides. After fixation, samples were permeabilized with 0.5% Triton X-100 in PBS for 15 min at room temperature. Samples were blocked with 3% BSA in PBS for 30 min at room temperature before incubation with primary antibody diluted 1:200 in blocking buffer (3% BSA, 0.05% Triton X-100 PBS) at 4 °C overnight in a humidified chamber. Samples were washed with PBS three times for 2 min each and then incubated with secondary antibody diluted 1:500 in blocking buffer at room temperature for 1 hr. Samples were washed with PBS and fixed with 4% PFA for 20 min. Fixed samples were further permeabilized with ice-cold 0.7% Triton X-100 per 0.1 M HCl (diluted in PBS) for 10 min on ice. DNA was denatured by 1.9 M HCl for 30 min at room temperature, followed by dehydration in ascending ethanol concentrations (70%, 85%, and 100% EtOH for 2 min each). Diluted FISH probes (probes and hybridization buffer from Empire Genomics) were separately denatured at 75 °C for 3 min and left to cool at room temperature for 5 min before adding onto glass slides covered with coverslips. After incubation at 37 °C overnight in a humidified chamber, samples were washed with 0.4× SSC, 0.1% Tween-20 in 2× SSC, and 2× SSC for 2 min each to get rid of non-specific binding, followed by DAPI (50 ng ml^−1^ diluted in 2× SSC or double-distilled H_2_O) staining for 2 min. Finally, samples were washed with 2× SSC and double-distilled H_2_O for 2 min each before air-drying and mounting with Prolong Diamond.

### Metaphase DNA-FISH

Cells were grown to 70% confluency in a 6cm dish and treated with KaryoMAX Colcemid (Gibco) for 4 hr. After a mitotic shake off, the media was collected. The remaining cells were treated with trypsin for 2 min. Fresh media was added to deactivate the trypsin, and the media was collected and centrifuged at 400g for 4 min. The cell pellet was resuspended in 600 µL of 75 mM KCl and incubated at 37 °C for 20 min. 600 µL of Carnoy’s solution (3:1 methanol: acetic acid) was added dropwise to the cell suspension and centrifuged at 3000 rpm for 2 min in a swing bucket centrifuge. The supernatant was discarded, and the cell pellet was resuspended in another 600 µL of Carnoy’s solution and centrifuged at 3000 rpm for 2 min. This wash step was repeated a total of 3 times. The final pellet was resuspended in 200 µL of Carnoy’s solution. 10 µL of the resuspended cell solution was dropped onto preheated glass slides from a height of approximately 15 cm and left to air-dry. The slides were dehydrated in ascending ethanol concentrations of 70%, 85%, and 100% for 2 min each. Diluted FISH probes (Empire Genomics) were added to the slides and covered with a glass coverslip. The samples were denatured at 75 °C for 3 min and hybridized at 37 °C overnight in a humidified slide moat. After overnight hybridization, the samples were washed with 0.4× SSC, 0.1% Tween-20 in 2× SSC, and 2× SSC for 2 min each, followed by DAPI staining for 2 min. Finally, samples were washed with 2× SSC and double-distilled H_2_O for 2 min each before air-drying and mounting with Prolong Diamond.

### Microscopy

Images were acquired on a Leica DMi8 widefield microscope using a 63× oil objective. Each *z* plane was post-processed by small-volume computational clearing on LAS X before generating maximum-projection images.

### IF and DNA FISH image analysis

Maximum-projection images of each sample were first imported into Aivia software (Leica Microsystems) as a stitched image. Individual machine-learning-based pixel classifiers were trained on the channels corresponding to the FISH probes of interest and DAPI to generate confidence masks for FISH foci and nuclei, respectively. The confidence masks were inputted into a recipe that assigned segmented FISH foci to segmented nuclei. Summed outline area of FISH foci was used to estimate ecDNA copy number per cell. IF signal, or abundance of proteins of interest, was measured using total (channel) pixel intensity per segmented nucleus or by foci area, as specified in the main text and figure legends.

### Metaphase DNA FISH image analysis

Colocalization analysis for two-color metaphase FISH data for ecDNA in TG19 ec cells described in Extended Data Fig. 2 was performed in Aivia. Individual machine-learning-based pixel classifiers were trained on the channels corresponding to the *MYC* and *tetO* FISH probes. The pixel classifiers were exported as confidence masks, which were used to create a recipe for segmenting out individual FISH foci. Object IDs and Nearest Object Distance measurements for segmented *MYC* and *tetO* FISH foci were exported from Aivia. *MYC* and *tetO* foci were classified as colocalized if the nearest object distance was less than or equal to 1 unit, and overlap fraction was calculated by taking the fraction of *MYC* foci colocalizing with *tetO* foci over total *MYC* foci in each metaphase spread.

### Engineering of isogenic TG19 ec and TG19 HSR cell lines

The TG19 ec cell line was engineered as described previously^48^. The TG19 HSR line was engineered similarly from COLO 320HSR cells obtained from ATCC and detailed as follows. First, CRISPR-Cas9 was used to knock in a 96× *tetO* array into the intergenic sites between *MYC* and *PCAT1*. The cells were treated with puromycin to select for *tetO*-positive cells. After selection, lentiviral infection of TetR-mNeonGreen was performed, and FACS was used to sort and collect mNeonGreen positive cells. Upon TetR-mNeonGreen binding to *tetO*, *tetO* foci fluoresce green. Monoclonal cell populations for both ecDNA(+) and HSR(+) cells were isolated via limiting dilution. Promising clones were evaluated by microscopy, and lentiviral infection of H2B-emiRFP670 was performed to label histone H2B protein in far red. FACS was used to sort and collect double-positive emiRFP670 and mNeonGreen cells. Another round of isolation of single clones by limiting dilution was performed to generate the final monoclonal TG19 ec (clone D11_2) and TG19 HSR (clone B2) cell lines that exhibited bright TetR-mNeonGreen foci with low background/noise. Metaphase DNA FISH was performed to evaluate *tetO* labeling efficiency of *MYC* and verify amplicon type remained unchanged.

### Live-cell imaging of ecDNA

10K cells were seeded in poly-D-lysine or fibronectin-coated wells of a 96-well glass bottom plate approximately 24 hours before imaging. Immediately before imaging, the medium was carefully replaced with FluoroBrite DMEM (Gibco, A1896701) supplemented with 10% FBS, 1× GlutaMAX, and 1:200 Prolong live antifade reagent (Thermo Fisher Scientific, no. P36975). Cells were imaged on a top stage incubator (Okolab) fitted onto a Leica DMi8 widefield microscope with a 63× oil objective. Temperature was maintained at 37 °C, and humidity and CO_2_ (5%) were controlled throughout the imaging time course.

### Live-cell image analysis: lineage tracking

Lineage tracking was performed manually by recording the time of cell division of the single initial progenitor and its subsequent progeny over the duration of image acquisition. Initial seeding density of 10K cells and acquisition timestep of 3 hr allowed individual divisions to be followed with high confidence.

### Live-cell image analysis: copy number quantification

Timelapse images were cropped by relevant timeframes (according to lineage tracking results) and exported as maximum-projection images in LAS X software. Images per timeframe were imported into Aivia. Outlines of nuclei of interest in each lineage were manually drawn, and Cell IDs were recorded. A machine-learning-based pixel classifier was trained on the TetR-mNeonGreen foci and exported as a confidence mask. The confidence mask was inputted into a recipe that assigned segmented FISH foci to the outlined nuclei. Summed outline area of the TetR-mNeonGreen foci was used to estimate ecDNA copy number per cell. Cells in which copy number (by TetR-mNeonGreen foci area) could not reliably be measured and estimated in the parent cell due to chromatin condensation at time of image acquisition were excluded from analysis.

### Bootstrapping LOESS minima

To compare the optimal copy number associated with lowest T2D between control and CHK1i-treated cells, we fit LOESS models to visualize the relationship between T2D and TetR-mNeonGreen foci in each group. We then used bootstrap resampling (1,000 iterations) to estimate the location of the minimum of the smoothed curve. The median and 95% bootstrap confidence interval were used to identify the optimal copy number in each group.

### Whole genome sequencing (WGS) of cell lines and xenograft tumour tissues

Genomic DNA from cultured cells in a confluent six-well dish was extracted using the QIAamp DNA Mini Kit (Qiagen) according to manufacturer’s protocols. Single cells were collected and resuspended in 200 μL PBS. 20 μL of proteinase K and 200 μL of Buffer AL were added and mixed thoroughly by pulse-vortexing for 15 s. Samples were incubated at 56 °C for 10 min. 200 μL of EtOH was added, and samples were pulse-vortexed for 15 s. The samples were then pipetted into a QIAamp Mini spin column and centrifuged at 6000g for 1 min. The flow-through and collection tube were discarded, and 500 μL of buffer AW1 was added to the spin column placed in a new collection tube. After centrifugation at 6000g for 1 min, the filtrate was discarded and 500 μL of buffer AW2 was added. Samples were centrifuged at 20000g for 3 min, and the filtrate was discarded. The spin column was then transferred to a new 1.5 mL microcentrifuge tube, and DNA was eluted by adding 200 μL buffer AE to the center of the spin column membrane. Samples were incubated at RT for 1 min before being centrifuged at 6000g for 1 min. This final step with buffer AE was repeated to increase the yield of DNA.

WGS library preparation was performed with the FS DNA Library Prep Kit from NEB according to the manufacturer’s protocol and with the following parameters: 250 ng gDNA used as input, fragmentation performed with incubation time of 15 min to yield 200 to 450 bp fragments, approximate final library size distribution of 320 to 470 bp (1st bead selection volume = 30 μL and 2nd bead selection volume = 15 μL), and final PCR amplification performed for four cycles. PE150 sequencing was performed on NovaSeq to yield at least 10X coverage at Novogene. Adaptor sequences and low-quality reads were removed from raw fastq files by Trimmomatic PE (version 0.39). Then, the cleaned fastq reads were aligned to hg38 and analyzed by AmpliconSuite-pipeline to call copy numbers and amplicon features (v1.3.5).

For xenograft tumours, genomic DNA was extracted from 10 to 30 mg of excised tumour tissue using the DNeasy Blood & Tissue Kit. Tissue was cut into small pieces using micro-dissecting scissors and placed in a 1.5 mL microcentrifuge tube. Then, 20 μL of proteinase K and 180 μL of buffer ATL were added. The mixture was pulse-vortexed for 15 s and incubated at 56 °C until completely lysed (approximately 3 hours, with pulse-vortexing for 15 s every 30 min). As described above, buffer AL and EtOH were added, and the mixtures were pipetted into DNeasy Mini spin columns. Centrifugation steps, with addition of buffers AW1, AW2, and finally AE were performed as described above. Library preparation (PCR+) and sequencing (Illumina – NovaSeq X Plus – PE150) to yield at least 10X coverage was performed at Novogene.

### Paired scATAC-seq and scRNA-seq library generation

Single-cell paired RNA-seq and ATAC-seq libraries were generated using the 10x Chromium Single Cell Multiome ATAC + Gene Expression platform according to the manufacturer’s protocol. Then, libraries of PE150 sequencing were performed on NovaSeq at Novogene. Data for COLO 320DM and COLO 320HSR cells were generated previously and published under BioProject Accession: PRJNA671462^11^. GBM39KT, SNU16, SNU16m1, TR14 cells were generated previously and published under BioProject Accession: PRJNA1127616^48^.

### Paired scATAC-seq and scRNA-seq analysis

Paired single-cell RNA-seq and ATAC-seq reads were aligned to the GRCh38 reference genome (refdata-gex-GRCh38-2020-A) using cellranger-arc count (10x Genomics, v2.0.2). Cells with more than 10,000 unique scATAC-seq fragments were retained for downstream analysis. Amplicon copy numbers were inferred by calculating fold changes between large genomic intervals and their local background using our previously developed and validated method^19,48^. To mitigate sparsity and peak-specific biases in scATAC-seq, we used 1-Mb windows with 200-kb sliding steps. Intervals overlapping ENCODE hg38 blacklist regions were removed due to their tendency to exhibit artificially high ATAC-seq signal. Focal amplicon regions identified by the AmpliconSuite pipeline were excluded from background estimation to avoid inflation of diploid regions by high-copy loci. For each window, we computed the average log2(fold change) relative to the 100 nearest background intervals; assuming a diploid baseline for most of the genome, copy number was calculated as 2 × 2^log2(fold change). The oncogene amplicon region was defined as the window containing the gene of interest with the highest inferred copy number and was added to the Seurat object for copy number–related analyses.

Aligned scRNA-seq matrices were processed using Seurat (v5.3.1). Cells with >200 detected genes, >500 total counts, and <30% mitochondrial RNA content were retained. RNA counts were log-normalized using NormalizeData and scaled using ScaleData. Gene set enrichment scores were computed with UCell (v2.14.0). Hallmark (H) and C2 (Curated) gene sets were obtained via the msigdbr package, and pathway activity scores were generated using UCell’s AddModuleScore_UCell and SmoothKNN functions.

### Fluorescence Activated Cell Sorting (FACS) of GBM39 cells

GBM39-EC and GBM39-HSR cells were prepared for FACS as described previously^14^. Briefly, cultured cells were treated with TrypLE and passed through a cell filter to make a single-cell suspension. The cells were suspended in flow cytometry buffer (Hanks’ Balanced Salt Solution buffer without calcium and magnesium, 1× GlutaMAX, 0.5% (v/v) FCS, 10 mM HEPES). EGFRvIII monoclonal antibody 806^47^ was added at 1 μg per million cells and incubated on ice for 1 h. Cells were washed in flow cytometry buffer and resuspended in buffer with anti-mouse Alexa Fluor 488 antibody (1:1,000, Thermo Fisher Scientific, no. A11017) for 30 min in the dark. Cells were washed and resuspended in flow cytometry buffer at approximately 5 million cells per milliliter. Cells were sorted using BD FACSAria™ II Flow Cytometer, which was calibrated; gating was informed using negative controls. Cells were sorted into the following three groups: High = top 10%, Mid = middle 15%, Low = bottom 10% according to Alexa Fluor 488 expression.

Sorted cells from each copy number population were maintained in culture and reserved for two downstream assays: 1) *EGFR* FISH and 2) cell counting. On days 0, 3, 5, and 7, 50K were fixed with 4% PFA and subsequently cytospun onto glass slides for subsequent *EGFR* FISH staining to assess changes in copy number over time. For cell counting, 3K cells from each group were seeded into one well of a 96-well round (U) bottom plate (3 wells per group per day for days 1, 2, and 3) and counted using a hemocytometer.

### GBM39-EC and GBM39-HSR cell counting after FACS

Cell counting was performed on days 1, 2, and 3 after seeding sorted GBM39-EC and GBM39-HSR cells in a 96-well round (U) bottom plate. 100 µL cell suspensions in each well were gently pipetted up and down three times to break up clumps before being transferred to a corresponding well containing 100 µL of TrypLE. The cells were gently pipetted up and down three times and incubated with TrypLE for 3 to 5 min. 100 µL of fresh media was added to deactivate TrypLE, and the plate was centrifuged for 4 min at 400g. The supernatant was carefully discarded, and the cell pellet was resuspended in 20 µL of fresh media. 10 µL of the resuspended cell solution was mixed with 10 µL of Trypan blue and loaded into the counting chamber of a hemocytometer. The hemocytometer was placed under a light microscope, and live and dead cells within the center and four corner squares were counted.

### Xenograft mouse models and tumourigenicity assay

Xenograft mouse model tumours were generated as described previously^37^. Briefly, female athymic nude mice (Charles River Laboratories) were housed under standard conditions. Immediately before subcutaneous cancer cell injection, mice were anesthetized using isoflurane in an induction chamber. A 100 µL aliquot of the prepared cell suspension (1:1 mixture of PBS: Matrigel) was injected into the subcutaneous space of the flanks bilaterally using a 25-gauge needle and a 1 mL syringe. For tumourigenicity assays, 125K COLO 320DM cells or 125K COLO 320HSR cells were injected into two groups of mice (n=5 in each group). Mice were monitored post-procedure for signs of distress or complications. Tumour length and width were measured several times per week until the experimental endpoint (days 20 for tumourigenicity assays, 28 for competition assay), at which time mice were euthanized using a sealed CO_₂_ chamber, with cervical dislocation performed as a secondary confirmatory method. Tumours were sharply dissected from the subcutaneous space, immediately flash-frozen in liquid nitrogen, and stored at −80 °C for future analysis.

### Simulation of cancer cell growth under optimal copy number model

Algorithms of three models of ecDNA copy number and T2D (**Extended Data Fig. 6a-c**) are: (1) a horizontal model: In this model, proliferative fitness is independent of ecDNA copy number. Where 𝒩(mean, sd) is a normal distribution, 𝑇_min_ enforces a minimum division time:

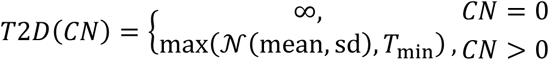

1. a sigmoid-plateau model: This model captures a dose-response-like increase in fitness with ecDNA copy number that plateaus at high copy numbers, without penalizing excess ecDNA abundance. Oncogenic benefit was modeled as a logistic function of ecDNA copy number:

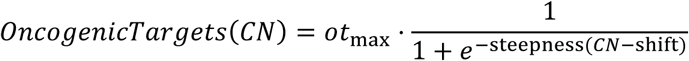

Total fitness was defined as the sum of a baseline term and oncogenic benefit, and cell doubling time (T2D) was assumed to be inversely proportional to fitness; where steepness=1.0, shift=5.0, 𝑜𝑡_max_=1.0, baseline=0.02:

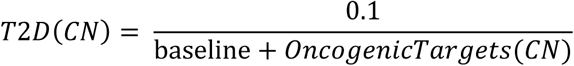

1. a quadratic model: This model represents a non-monotonic fitness landscape derived from live-cell imaging data, with maximal fitness at an intermediate copy number. Where coefficients a=0.2978, b=-5.4405, c=55.5155 fitted from experimental measurements, and the minimum of the parabola defines the optimal ecDNA copy number range:

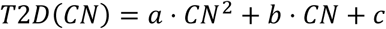

EcDNA(+) and HSR(+) cell population growth was simulated by modeling the time to division (T2D) as a quadratic function of amplicon copy number, with parameters empirically derived from live-cell imaging. Using this relationship, we defined the function add_ct to generate a matrix of amplicon copy numbers and their corresponding T2D values. For ecDNA(+) cells, copy number inheritance was simulated using a binomial distribution to model the uneven segregation of ecDNA during mitosis. The function amp_random_inheritance uses the matrix of copy numbers and T2Ds, together with parameters governing random inheritance, to simulate ecDNA(+) population growth over a specified number of cell cycles. To model HSR(+) populations, we defined amp_fixed_inheritance, in which amplicon copy numbers were divided equally between daughter cells, reflecting the fixed chromosomal inheritance of HSRs.

### Engineering of COLO 320DM-mCherry and COLO 320HSR-GFP reporter cell lines

pLV-mCherry (Addgene plasmid no. 36084) and pLV-eGFP (Addgene plasmid no. 36083) were gifts from Pantelis Tsoulfas^49^. Plasmids were packaged and transfected to produce lentivirus according to Addgene Lentivirus Production protocol. Viruses were harvested and stored at −80 °C. COLO 320DM cells were infected with mCherry-packaged lentivirus (mixed with 1:1000 10 mg/mL polybrene in fresh media), and COLO 320HSR cells were similarly infected with eGFP-packaged lentivirus. After two days, cells were treated with ampicillin to select for successfully transformed cells.

### In vitro competition assay

400K COLO 320DM-mCherry and COLO 320HSR-GFP cells were co-cultured at each of the following ratios: 100% DM, 75% DM: 25% HSR, 50% DM: 50% HSR, 25% DM: 75% HSR, and 100% HSR. 48 hr after plating, cells were collected, spun down 2x at 400g for 5 min and stained with live-dead dye. To label dead cells, cells were stained with LIVE/DEAD Fixable Far Red Dead Cell Stain (Thermo Fisher Scientific, no. L34973) or Sytox Blue Live/Dead Cell Stain (Thermo Fisher Scientific, no. S34857). Flow cytometry was performed to quantify proportion of each cell type.

### In vivo competition assay

120K COLO 320DM-mCherry and COLO 320HSR-GFP cells were injected into mice at each of the following ratios: 100% DM, 75% DM: 25% HSR, 50% DM: 50% HSR, 25% DM: 75% HSR, and 100% HSR. Tumours were excised at 28 days and dissociated according to 10x Genomics Tissue Fixation and Dissociation Demonstrated Protocol #CG000553 (Rev B). After staining with live-dead dye, flow cytometry was performed to quantify proportion of each cell type.

### Flow cytometry

For in-vitro assays, COLO320DM and HSR cell lines were seeded at 400,000 cells/mL in full media (DMEM with 10% fetal bovine serum). Upon plating, cells were subsequently trypsinized, collected, washed, and fixed in 4% paraformaldehyde (PFA) in PBS. For in-vivo assays, resected tumour sections were homogenized into cell suspensions, upon which tumourigenicity was quantified using flow cytometry. For both in-vivo and in-vitro assays, the extent of proliferation was quantified using the Symphony analyzers at the Stanford Shared Fluorescence Activated Cell Sorting (FACS) facility using a NIH S10 Shared Instrument Grant (Symphony 1S10OD026831-01). Flow cytometry data was analyzed using FlowJo software (BD). Gating strategy was established from single-reporter populations.

### DNA FISH and IF staining on FFPE tissue sections

Excised tumour tissues were sectioned onto glass slides by the Stanford Human Pathology/Histology Service Center. Samples were first dewaxed by incubating slides upright at 65 °C for at least 1 hr. Slides were then submerged in xylene for 10 min 2x. Tissue re-hydration was performed by immersing slides in descending EtOH gradations as follows: 100%, 5 min, 2x; 95%, 3 min, 2x; 80%, 3 min, 1x, 70%, 3 min, 1x, 50%, 3 min, 1x. The slides were rinsed in double-distilled H_2_O before proceeding with antigen retrieval. Slides were immersed in 1× Tris-EDTA Antigen Retrieval Buffer (10× Tris-EDTA Antigen Retrieval Buffer made by combining 10 mM Tris Base, 1 mM EDTA solution, 0.05% Tween-20) at 90-95 °C for 15 min. After cooling, dehydration was performed by immersing slides in ascending EtOH gradations as follows: 50% EtOH, 3 min, 1x; 70% EtOH, 3 min, 1x; 80% EtOH, 3 min, 1x; 95% EtOH, 3 min, 1x; 100% EtOH, 3 min, 1x and left to to air-dry. *MYC* FISH probe (Empire Genomics) was diluted in hybridization buffer and pre-annealed at 63 °C for 5 min. The probe was then applied to the samples, and the sections were covered with coverslips. The samples were denatured at 75 °C for 5 min and hybridized overnight at 37 °C in a humidified slide moat.

Following overnight hybridization, slides were added to warm 0.4× SSC/0.3% Tween-20. Coverslips were carefully removed, and slides were equilibrated in 2× SSC for 5 min. Slides were washed in 0.4× SSC/0.3% Tween-20 for 2 min, with vigorous shaking for the first 10 to 15 sec. Slides were washed in 2× SSC/0.1% Tween-20 for 5 min and then rinsed in double-distilled H_2_O for 5 min. Slides were fixed with 4% PFA for 10 min and washed with PBS for 5 min. Slides were lightly permeabilized with 0.025% Triton X-PBS for 10 min. Slides were then washed in PBS for 5 min 3x. Blocking was performed with 3% goat serum for at least 1 hr, and Ki67 primary antibody diluted (1:200) in blocking buffer was applied. Slides were incubated with primary antibody overnight at 4 °C in a humidified chamber.

After overnight incubation, slides were washed with PBS for 5 min 3x. Secondary antibody diluted (1:500) in blocking buffer was applied, and slides were incubated with secondary antibody for 1 to 2 hr at room temperature. Samples were then washed with PBS for 5 min 3x and then treated with TrueView Autofluorescence Quenching Kit for 2 min. DAPI was applied for 5 min, and slides were washed with 2× SSC and double-distilled H_2_O. Finally, slides were air-dried before mounting with Prolong Diamond. Unless otherwise specified, all wash steps were performed at room temperature.

### Quantification and statistical analysis

Plots were prepared using R language for statistical computing. Unless otherwise specified, statistical tests were two-sided. Details of statistical tests can be found in the main text, figure legends, and Methods.

### Data availability

Single-cell multiomics (10x Chromium Single Cell Multiome ATAC + Gene Expression) data of COLO320DM and COLO320HSR were generated previously and published at the GEO (GSE159986)^11^. Single-cell multiomics data of SNU16m1, GBM39KT, and TR14 were generated previously and published at NCBI SRA under BioProject accession PRJNA1127616^48^.

